# Modulating Hydrogel Stiffness Through Light-Based 3D Printing to Mimic Cardiac Fibrosis and Cardiomyocyte Dysfunction Using hiPSC-Derived Cells

**DOI:** 10.1101/2025.05.20.655137

**Authors:** Sogu Sohn, Nima Momtahan, Lynn M. Stevens, Jiwan Han, Yutong Liu, Meghan T. Kiker, Elizabeth A. Recker, Zachariah A. Page, Janet Zoldan

## Abstract

The human heart’s limited regenerative capacity is a significant barrier to addressing cardiovascular disease (CVD). This is particularly true for cardiac fibrosis, a form of CVD wherein the wound healing process has gone awry. In cardiac fibrosis, excessive scar tissue formation due to dysregulated remodeling of the heart’s extracellular matrix (ECM) results in increased stiffness that reduces cardiac output and can lead to heart failure. This dysregulated ECM deposition is driven by activated cardiac fibroblasts, where cell substrate stiffness is known to play a role in cardiac fibroblast activation. New preclinical models that accurately recapitulate the behavior of activated cardiac fibroblasts are needed to better understand and treat cardiac fibrosis. To this end, we describe a model wherein human induced pluripotent stem cell (hiPSC)-derived cardiac fibroblasts (HCFs) are cultured on 3D printed hydrogels of tunable stiffness, fabricated using dosage controlled digital light processing (DLP). We demonstrate that our model can induce HCF activation in the absence of TGFβ, a key mediator of fibroblast activation, surpassing the activation levels seen with HCFs activated with TGFβ on protein-coated tissue culture plates. Furthermore, combining stiffer hydrogels with TGFβ recapitulates fibroblast activation similar to what is observed in native cardiac tissue. Lastly, by indirectly coculturing HCFs seeded and activated on these stiff hydrogels with hiPSC-derived cardiomyocytes, we demonstrate that the activated HCFs in our cardiac fibrosis model can impair cardiomyocyte function, mimicking the deleterious effects of cardiac fibrosis.

## INTRODUCTION

Cardiovascular disease (CVD) is a leading cause of death with projections for growth, making it a prominent global public health concern.^1,2^ Contemporary intervention strategies to curb the progression of CVD include lifestyle changes, pharmaceuticals, and surgery, yet severe lasting tissue damage persists.^3,4^ The lack of effective treatments stems in part from our limited understanding of the complex mechanisms that lead to CVD, along with an inability to effectively mimic the complex extracellular matrix (ECM) environment necessary to test new treatments in vitro. It is known that an adult heart has limited regenerative capacity to manage CVD, forming non-functional scar tissue in response to damage to healthy tissue.^5–7^ This scar tissue is formed as part of the normal wound healing response to maintain heart chamber structural integrity and transmission of contractile forces.^5–7^ However, the increased stiffness of this scar tissue relative to healthy cardiac tissue, combined with the scar tissue’s inability to actively contract, makes it an inadequate replacement.^7^ Exacerbating the issue is the poorly regulated ECM deposition, which leads to excessive and continuous ECM build-up during wound healing in the heart, ultimately resulting in cardiac fibrosis, a common form of CVD.^7,8^ Critically, the accumulation of stiff and non-contractile fibrotic scar tissue during cardiac fibrosis leads to pathological thickening of heart structures, increasing the risk of arrhythmia and heart failure.^9^ Moreover, cardiac fibrosis can occur in conjunction with other forms of CVD and may persist even after the underlying causes of cardiac injury are resolved.^9^ Accordingly, accurate models of cardiac fibrosis are needed to better understand and treat it.

Preclinical models that recapitulate the target disease state represent a valuable approach to developing intervention strategies. Concerning preclinical models of CVD, research has focused on the use of cardiomyocytes (CMs) due to their principal role in cardiac function.^10,11^ However, mimicking the behavior of cardiac fibroblasts (CFs) to emulate cardiac fibrosis is key as CFs are non-myocyte cells in the cardiac niche that play vital roles in the heart’s development, normal function, and pathological response.^12–15^ CFs contribute to regulating the behavior of other cells in the cardiac niche, most notably CMs, through paracrine signaling. However, they are primarily known for regulating cardiac structure via secretion and degradation of cardiac ECM components.^12–14,16^ Notably, CFs promote maturation of the embryonic heart by mediating changes in cardiac ECM structure and CM behavior, leading to more compact and functionally mature cardiac tissue.^13,16^ In the normal postnatal heart, CFs regulate ECM synthesis and degradation to maintain tissue mechanics, facilitate electrical signal conduction to promote synchronous contractions, and secrete cytokines to influence cell behavior through both paracrine and autocrine signaling.^15^ In a stressed and damaged heart, CFs shift from a quiescent to an activated phenotype state, participating in the pathological remodeling of cardiac tissue and driving the development of cardiac fibrosis.^9,14^ These activated CFs (also known as myofibroblasts) display pro-inflammatory behavior characterized by increased proliferation and excessive ECM synthesis, disrupting the balance maintained by quiescent CFs.^9,17^ Furthermore, the increased stiffness of the scar tissue relative to the healthy cardiac tissue promotes CF activation, causing a snowball effect that raises cardiac fibrosis-associated risks.^15^ Thus, increased tissue stiffness is both a cause and effect of CF activation. Accordingly, preclinical models of cardiac fibrosis wherein CFs are cultured on a substrate with tunable stiffness are needed to mimic the heart’s stiffening as injured cardiac tissue is replaced with scar tissue.

Three-dimensional (3D) bioprinting has emerged as a strategy to enable the fabrication of biomimetic scaffolds that better recapitulate in vivo conditions. Bioprinting of hydrogels – water-swollen polymer networks – has successfully been employed to create tissue-like constructs, including cartilage, bone, and muscle.^18–20^ Notably, digital light processing (DLP) 3D printing offers a compelling combination of rapid fabrication speeds (>25 mm/hour), high microscale feature resolution (≤100 µm), minimal application of mechanical stress that supports cell encapsulation, and broad material compatibility.^21^ These unique features are accomplished through precise spatiotemporally controlled photocuring processes, where patterned light from a digital micromirror device is projected into a liquid resin vat containing monomers and a photoinitiation system. While DLP bioprinting has been used to develop artificial tissue models, the ability to pattern stiffness into hydrogels to systematically and accurately model cardiac fibrosis represents an unmet challenge.

Herein, we employ norbornene-functionalized gelatin (NorGel) and grayscale (i.e., dosage-controlled) projections to precisely modulate photocrosslinking, thereby tuning the stiffness of hydrogels designed for use as cell scaffolds. NorGel was chosen due to its biocompatibility, enzymatic degradability, and arginine-glycine-aspartic acid (RGD) motifs that promote integrin-mediated cell adhesion. Unlike methacrylate-functionalized gelatin (GelMA), which polymerizes via a free-radical chain-growth mechanism, NorGel undergoes a step-growth mechanism with thiol crosslinkers, facilitating temporal control over crosslink density. By varying light intensity for a fixed exposure time, hydrogel stiffness could be tuned across a range that recapitulates the foldchange increase in stiffness (2-10×) that healthy cardiac tissue (7-10 kPa) experiences when it becomes fibrotic (20-100 kPa).^22–24^ Seeding quiescent CFs derived from hiPSCs onto the 3D-printed hydrogels was shown to promote an activated CF phenotype. Moreover, by varying the hydrogel stiffness and dosing the hiPSC-CFs (HCFs) with drugs used to treat cardiac fibrosis, we demonstrated the model’s ability to induce changes in HCF gene expression that recapitulate key features of cardiac fibrosis, such as impaired cardiomyocyte function (**Fig. 1**).

**Figure 1.**
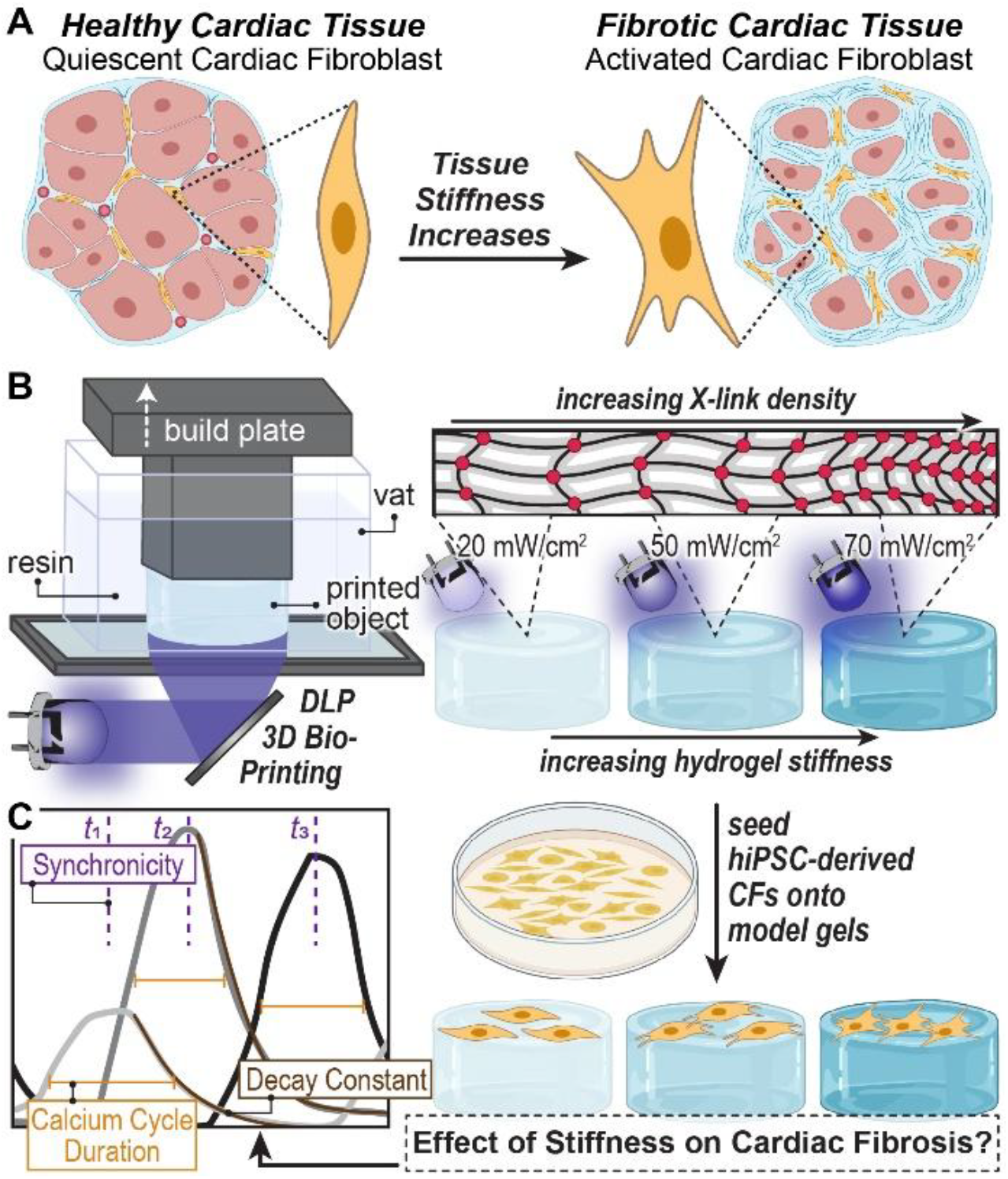
Modulating Hydrogel Stiffness via Light-Based 3D Printing to Study Cardiac Fibroblast Activation. A) Illustration of healthy vs. fibrotic cardiac tissue highlighting the transition of quiescent cardiac fibroblasts to activated fibroblasts in response to increased tissue stiffness (made with BioRender.com). (B) Digital Light Processing (DLP) 3D printing is used to fabricate hydrogels at varying light intensities (20, 50, 70 mW/cm^2^), modulating crosslink density and thereby hydrogel stiffness. C) Human induced pluripotent stem cell (hiPSC)-derived cardiac fibroblasts seeded onto the 3D printed hydrogels were used to correlate hydrogel stiffness to cardiomyocyte dysfunction via electrochemical coupling metrics (made with BioRender.com).

## METHODS

### Materials

#### Reagents

All chemicals were used as received unless otherwise noted. (2,2,6,6-tetramethylpiperidine-1-yl)oxidanyl (TEMPO, 98%), 5-norbornene-2-carboxylic acid (mixture of *endo* and *exo*, predominantly *endo*, 98%), and gelatin from bovine skin (Type B) were purchased from Sigma-Aldrich. 1,4-Dithio-DL-threitol (DTT, ≥99%, Assay by titration) and *N*-hydroxysuccinimide (NHS) were purchased from Chem Impex. 1-(3-Dimethylaminopropyl)-3-ethylcarbodiimide hydrochloride (EDC) and lithium phenyl(2,4,6-trimethylbenzoyl)phosphinate (97%) (LAP) were purchased from Ambeed. Cell Culture Phosphate Buffered Saline (PBS) 1X without calcium and magnesium was purchased from Corning. Deuterated water (D_2_O) (99.9 atom % D, contains 0.05 wt. % 3-(trimethylsilyl)propionic-*2,2,3,3-d4* acid, sodium salt) was purchased from Sigma Aldrich (product nm. 450510).

### NorGel Synthesis and Characterization

#### Synthesis

Norbornene functionalized gelatin (NorGel) was prepared using a modified procedure reported by Van Hoorick et al. 2018^25^ (Figure S1). 5-norbornene-2-carboxylic acid (2.0 g, 14.5 mmol) was dissolved in 50 mL dry dimethyl sulfoxide (DMSO) under inert conditions (N_2_). After complete dissolution, 2.5 equivalents of EDC (1.86 g, 9.7 mmol) (relative to the primary amines present in 10 g gelatin, i.e., 0.38 mmol/g gelatin) and 1.5 equivalents of NHS (relative to EDC, 1.67 g, 14.55 mmol) were added and stirred at room temperature under inert conditions for 25 hours. In a separate flask, 10 g of Type B gelatin was dissolved in 150 mL dry DMSO. The mixture was heated to 50 °C under inert conditions (N_2_). Following the complete dissolution of the gelatin, the 5-norbornene-2-succimidylester mixture was transferred to the gelatin solution via cannulation. The mixture was stirred at 50 °C under inert conditions for 20 hours. Next, the mixture was precipitated into acetone and filtered. The precipitate was dissolved in deionized water and dialyzed (Sigma Aldrich cellulose membrane MWCO ∼14000, D9402-100FT) for 24 h at 45 °C with a minimum of 5× water exchanges. After dialysis, the pH was adjusted to 7.4 via the dropwise addition of a 1M NaOH_(aq)_ solution, as monitored using pH test strips (Millipore Sigma, Cat # M1095350001). In a final step, the aqueous NorGel solution was lyophilized to remove water and obtain the product as an off-white solid (∼90% yield). ^1^H NMR (400 MHz, D_2_O) *δ* 7.32-7.35 (m, gelatin backbone), 6.22 – 5.90 (m, norbornene double bond, 2H), 4.6 – 0.91 (m, gelatin backbone), 0.0 (s, TMSP reference), see Figure S2 in the supporting information for additional details.

#### Quantification of Norbornene Functionalization

NorGel was dissolved in deuterated water (D_2_O, Sigma Aldrich product no. 450510) at a known weight fraction. The D_2_O contained 0.05 wt% 3-(trimethylsilyl)propionic-2,2,3,3-d4 acid and sodium salt (TMSP), which was used as an internal reference. ^1^H NMR spectra were collected and analyzed using Mestrenova software. TMSP was referenced to a chemical shift of 0.0 ppm, the phase was corrected manually, and the baseline was corrected using a polynomial fit (polynomial order set to 3). The norbornene C=C–H proton signals located at 6.22-5.90 ppm were integrated and set to 2 (the number of corresponding hydrogen atoms), after which the TMSP signal located at 0.00 ppm could be integrated (Figure S2). Using Equation S1 in the supporting information, the moles of norbornene per gram of NorGel could be determined. Following the calculation of moles of norbornene per gram of NorGel, a stoichiometric amount of dithiothreitol (DTT) by norbornene to thiol functionality (*f* = 2 for DTT) was determined using Equation S2 in the supporting information (Table S1).

### Resin Formulation and 3D Printing

#### Formulation

NorGel resins were prepared by dissolving NorGel in PBS (20 wt% NorGel in 80 wt% PBS), followed by the addition of photoinitiator, lithium phenyl-2,4,6-trimethylbenzoylphosphinate (LAP; 0.3 wt% relative to bulk), and radical inhibitor, 2,2,6,6-Tetramethylpiperidin-1-oxyl. (TEMPO; 0.1 wt% relative to bulk). After mixing with mild heat (<40 °C) in the absence of light to create a homogeneous solution, crosslinker DTT (0.2 wt% relative to bulk) was added and vortexed to dissolve.

#### DLP 3D Printing

3D printing was performed using a custom-made, digital light processing (DLP) 3D printer (MONO3MZ2, Monoprinter). The printer contained an LED projector with a wavelength centered at 405 nm (PDC04-405 nm). The LED intensity at the image plane was varied between 20 mW/cm^2^, 50 mW/cm^2^, and 70 mW/cm^2^. The projected image was a rectangle with 20 mm × 40 mm × 2 mm dimensions. A transparent fluorinated ethylene propylene (FEP) polymer film (Teflon FEP film, 127 μm thick, DuPont) was used as the base of the resin vat to provide a non-stick bottom surface. The resin vat was equipped with an ITO-coated glass bottom and a metal holder that was heated to 37 °C. Additional information about the printer and printing setting can be found in the Supplementary Methods section.

#### Post-printing Processing for Cell Culture

3D printed rectangles were submerged in PBS at 37 °C in the dark on a heater shaker set to 100 rpm overnight to fully swell and leech out any unreacted components (e.g., photoinitiator and monomer). Following swelling, 8 mm disks were cut out using a biopsy punch. Hydrogel disks were sterilized in 70% ethanol overnight at room temperature with gentle agitation on an orbital shaker. After the ethanol wash, samples were transferred to a biosafety cabinet and rinsed five times with Dulbecco’s phosphate-buffered saline (DPBS) without calcium and magnesium (Corning, 21-031-CM). Rinsed samples were placed in 12-well clear plates (one sample per well) with 1 mL Fibroblast Growth Medium 3 (FGM-3; PromoCell, C-23025) overnight in a cell incubator to assess sterility (i.e., whether culture resulted in the growth of biological contaminants).

### Rheology

#### In situ Photorheology

Photorheology was conducted using a Discovery Hybrid Rheometer from TA Instruments (DHR 20). The rheometer was equipped with a “UV Light Guide” accessory (part # 546301.901, TA Instruments), a 20 mm diameter disposable acrylic bottom plate (part # 403064.901, TA Instruments), and an HR x0 Upper Peltier Plate System (part # 534050.901, TA Instruments) containing an HR x0 Stainless Steel 20 mm (part # 534519.94) upper geometry. A liquid light guide was used to illuminate the samples with the corresponding visible light from a 405 nm LED (LCS-0405-12-22, Mightex Systems) connected to a driver from which light intensity could be controlled remotely (SLC-MA02-U, Mightex Systems). The rheometer was set to perform a single data acquisition cycle using the following experimental parameters: fast oscillation step at 1% strain and 1 Hz frequency for a total run time of 300 s. The gap height was set to 100 μm for each experiment to match 3D printing conditions. Photorheology was performed at 37 °C under air for all samples. The light was activated 10 seconds into the fast oscillation step. The storage modulus (*G′*) and the loss modulus (*G″*) were monitored in real-time, with samples run for 30s to determine the gel point or 60s to assess the plateau modulus. The crossover of *G′* and *G″* served as an estimate for the gel point, determined using the “moduli cross” data analysis function in TRIOS software from TA instruments. Each sample was tested six times at the printer light intensities. Gel point values were obtained by averaging all six trials, subtracting the initial 10 seconds corresponding to the light-off phase of the run.

#### Bulk Stiffness Measurements

Bulk hydrogel stiffness was characterized by measuring the storage modulus (G′) using a TA Instruments Discovery Hybrid Rheometer (DHR-20) following post-print washing. An 8 mm parallel steel plate head (part # 534515.941, TA Instruments) was used with the HR x0 Upper Peltier Plate System (part # 534050.901, TA Instruments). At 37 °C, a strain sweep was first conducted with a frequency of 10 rad/s and axial force of 0.5 N, followed by a frequency sweep at 1% strain in the linear-viscoelastic region, varying frequency from 0.1 rad/s to 100 rad/s.

### hiPSC Culture

hiPSCs were cultured in Essential 8 (E8) medium (Thermo Fisher, A1517001), with cells cultured as small colonies on tissue culture plates coated with recombinant human vitronectin (Thermo Fisher, A14700). E8 was exchanged daily until hiPSCs were approximately 70-85% confluent, at which point cells would be passaged. For single-cell passaging, hiPSCs were detached with Accutase (Accutase, AT104-500) and then centrifuged at 300RCF for 5 minutes, after which the cells were resuspended in E8 with 10 µM Y-27632 (Selleck Chemicals, S1049), and then counted using a hemacytometer. For maintenance culture, hiPSCs were seeded at 22,000 cells/cm^2^ in E8 with 10 µM Y-27632. For colony passaging, hiPSCs were suspended as small colonies by rinsing wells with DPBS and then incubating cells in 0.5 mM ethylenediaminetetraacetic acid (EDTA, Thermo Fisher, AM9260G). EDTA solution was then removed, and cells were gently sheared using E8 with a P1000 micropipette tip to generate a cell suspension of small colonies and reseeded at a 1:5 split.

### Quiescent Cardiac Fibroblast Differentiation

hiPSCs (WiCell, iPS DF19-9-11T.H) were differentiated into quiescent HCFs according to the protocol published by Zhang et al. in 2022^26^. Two days prior to the onset of differentiation, hiPSCs were seeded onto tissue culture plates coated with a 0.225 mg/mL solution of Geltrex (Thermo Fisher, A1413202) at a seeding density of 20,000 cells/cm^2^. Cells were cultured in Essential 8 (E8) medium (Thermo Fisher) supplemented with 10 µM Y-27632, with daily media exchanges using E8 without Y-27632. To initiate differentiation (Day 0), the medium was replaced with RPMI 1640 (Thermo Fisher, 11875135) containing B27 Supplement without insulin (Thermo Fisher, A1895601; RB-) and 6 µM CHIR99021 (Biorbyt, orb762913). On day 2, the medium was replaced with RB-only. After an additional 24 hours, the medium was replaced with RB-supplemented with 5 µM IWR-1 (Selleck Chemicals, S7086). On day 5, the medium was replaced with RB-only. After an additional 24 hours, cells were disassociated using Accutase and replated onto Geltrex-coated tissue culture plates at a seeding density of 20,000 cells/cm^2^. Cells were cultured in Advanced DMEM (Thermo Fisher, 12634028) and GlutaMax (Thermo Fisher, 35050061) mixture (ADGM) supplemented with 5 µM CHIR99021 and 2 µM retinoic acid (Sigma Aldrich, R2625). On day 7, the medium was exchanged for ADGM supplemented with 5 µM CHIR99021 and 2 µM retinoic acid. On day 9, CHIR99021 and retinoic acid were removed. On day 11, cells were disassociated with Accutase and replated onto Geltrex-coated plates at a split of 1:3-1:6 and cultured in ADGM supplemented with 2 µM SB431542 (Selleck Chemicals, S106710MM/1ML). On day 14, cells were disassociated again with Accutase and replated onto Geltrex-coated plates at a seeding density of 10,000 cells/cm^2^. Cells were cultured in Fibroblast Growth Medium 3 (FGM-3; PromoCell, C-23025) supplemented with 20 ng/mL fibroblast growth factor 2 (FGF-2; R&D Systems, 233-FB-010) and 10 µM SB431542 with media exchanges every other day. From day 20 onwards, cells were maintained in FGM-3 supplemented with 10 µM SB431542, exchanged every 48 hours. The HCFs used in subsequent experiments were expanded and passaged from cells from day 20 of differentiation.

### Immunostaining for Fluorescence Microscopy

Cells were rinsed twice with DPBS before incubating in 4% paraformaldehyde (PFA; Polysciences, 00380-1) at room temperature for 10 minutes. PFA was removed, and cells went through two rounds of 5-minute room-temperature incubations in DPBS with 300 mM glycine to neutralize the remaining PFA. Cells were then permeabilized in DPBS with 0.2% Triton X-100 for 10 minutes at room temperature before being washed with DPBS. Non-specific antigens were then blocked via incubating cells in a blocking buffer (DPBS with 1% bovine serum albumin and 0.1% Tween-20) for 30 minutes at room temperature. Cells were then incubated with primary antibodies diluted in blocking buffer at 1:200 dilutions overnight at 4 °C, with vimentin (Invitrogen, MA5-14564) and alpha-smooth muscle actin (αSMA; Invitrogen, ab5694) primary antibodies used to target HCFs and activated HCFs, respectively. Cells were then rinsed twice with blocking buffer before incubating with 1:1000 diluted fluorophore-conjugated secondary antibodies in blocking buffer for 1 hour at room temperature, with anti-rabbit AF594 (Abcam, ab150076) and anti-mouse AF488 (Invitrogen, A-11001) secondary antibodies used to label HCFs and activated HCFs, respectively. Cell nuclei were then stained using a 1:10,000 dilution of 4′,6-diamidino-2-phenylindole (DAPI) in blocking buffer for 1-2 minutes and then rinsed twice with DPBS. Fixed and stained samples were protected from light and stored at 4 °C until imaging. Additional information on primary and secondary antibodies used can be found in the supporting information (Table S2).

### Immunostaining for Flow Cytometry

Cells were dissociated with Accutase and centrifuged at 300 RCF for 5 minutes. The cell pellet was resuspended in 4% PFA and incubated on ice for 10 minutes. PFA was neutralized by diluting the cell suspension with DPBS containing 300 mM glycine, pelleting the cells at 300 RCF for 5 minutes, and resuspending the pellet in 300 mM glycine before pelleting once more. Cells were resuspended in DPBS containing 0.2% Triton and kept on ice for 10 minutes to permeabilize cell membranes. Permeabilized cells were washed in DPBS and then pelleted. Cells were resuspended in blocking buffer and incubated for 30 minutes on ice with gentle agitation on an orbital shaker. A fraction of the blocked cell suspension was aliquoted to keep as a negative control and not exposed to primary antibodies. The remaining cell suspension was incubated on ice in 1:100 dilutions of primary antibody solutions with gentle agitation on an orbital shaker for 1 hour, with vimentin (Invitrogen, MA5-14564) and alpha-smooth muscle actin (αSMA; Invitrogen, ab5694) primary antibodies used to target HCFs and activated HCFs, respectively. Cells were then pelleted/washed twice with blocking buffer before being resuspended in fluorophore-conjugated 1:500 dilutions of secondary antibody solutions and incubated on ice with gentle agitation on an orbital shaker for 30 minutes, with anti-rabbit AF594 (Abcam, ab150076) and anti-mouse AF488 (Invitrogen, A-11001) secondary antibodies used to label HCFs and activated HCFs, respectively. Negative controls were also stained with secondary antibodies to account for any non-specific binding. Cells were then pelleted/washed in sorting buffer (DPBS with 0.5% bovine serum albumin and 2mM EDTA) twice before being strained through 40 µm mesh caps into 5 mL sorting tubes. Additional information on primary and secondary antibodies used can be found in the supporting information (Table S2).

### Quantifying Relative Gene Expression

#### RNA Isolation

RNA was isolated from day 20 HCFs and hiPSCs (day 0 of differentiation) using an RNeasy Mini Kit (Qiagen, 74104). Isolated total RNA was eluted using nuclease-free water, with purity and concentration quantified using a Cytation 3 plate reader (BioTek Instruments).

#### Reverse Transcription

Total RNA was reverse transcribed into cDNA using a High-Capacity cDNA Reverse Transcription Kit (Applied Biosystems, 4368814). The resulting cDNA was kept on ice for immediate use in the qPCR reaction or stored at -80 °C.

#### qPCR

PCR amplifications were performed with PrimeTime qPCR Primers (Integrated DNA Technologies) and PowerUp SYBR Green (Thermo Fisher, A25742) as an intercalating dye. Relative gene expression was calculated using the ΔΔC_T_ approach, with GAPDH used as the housekeeping gene for all experiments. Primer assay IDs and sequences are available in supporting information (Table S3).

### Evaluating hiPSC-Cardiac Fibroblast Behavior

#### Characterizing HCFs

Day 20 hiPSC-CFs were passaged onto Geltrex-coated plates at 20,000 cells/cm^2^ and cultured in FGM-3 media supplemented with 10 µM SB431542. Media was refreshed daily until approximately 80% confluency was reached. Cells were then immunostained for downstream analysis using fluorescence microscopy or flow cytometry using previously described methods. For microscopy, vimentin and alpha-smooth muscle actin (αSMA) primary antibodies were used as markers for total HCFs and activated HCFs, respectively. For flow cytometry, HCFs were prepared using the previously described methods with a vimentin primary antibody to quantify total HCF yield and αSMA to quantify relative activated HCF yield. Relative changes in gene expression were assessed via qPCR for genes COL3A1, TCF21, and TBX20. Changes in the expression of these genes in HCFs were evaluated using day 0, undifferentiated hiPSCs as negative controls for reference.

#### Activating hiPSC-CFs on Tissue Culture Plates

To activate quiescent HCFs, the culture medium was exchanged for FGM-3 with transforming growth factor beta (TGFβ), a known promoter of CF activation, at concentrations of 0.1 ng/mL and 5 ng/mL for 48 hours. Because SB431542 inhibits the TGFβ signaling pathway and is part of the HCF maintenance medium, we used control samples of HCFs cultured in FGM-3 both with and without SB431542 to assess HCF activation, as TGFβ was added to FGM-3 media without SB431542. Changes in the activation of HCFs were evaluated qualitatively using fluorescence imaging and quantitatively using flow cytometry after staining samples with vimentin and αSMA to quantify changes in the proportion of activated HCFs. Changes in the relative expression of ACTA2, COL1A1, and COL3A1 in both control HCFs and HCFs exposed to 0.1 ng/mL TGFβ were measured to assess the expression of genes associated with the activated phenotype of HCFs.

#### Activating hiPSC-CFs on 3D Printed Hydrogel Disks

NorGel constructs were generated by 3D printing with photocrosslinking intensities of 20, 50, and 70 mW/cm^2^ to vary stiffness. The hydrogel disks were then washed with ethanol for sterilization, rinsed in DPBS, and allowed to air-dry before being fitted in 12-well clear tissue culture plates (Genesee Scientific, 25-106). Day 20 HCFs on Geltrex-coated tissue culture plates were disassociated using Accutase, pelleted, and resuspended in a single cell suspension in FGM-3 containing 10 µM SB431542. Cells were seeded at 30,000 HCFs per well onto hydrogel disks precisely cut to fill each well’s bottom surface, ensuring that cells adhered exclusively to the disk. The medium was exchanged after 24 hours to allow cells sufficient time to adhere to the hydrogels, with FGM-3 supplemented with 10 µM SB431542 or FGM-3 with 0.1 ng/mL TGFβ. Cultures were maintained under these conditions for 48 hours, after which cells were harvested for analysis. Fibroblast activation was evaluated by fluorescence imaging and flow cytometry following vimentin and αSMA staining, and by qPCR quantification of ACTA2, COL1A1, and COL3A1 expression. For flow cytometry, we determined both the proportion of αSMA positive cells and the degree of activation based on αSMA fluorescence intensity: events greater than 5,000 a.u. were classified as “high” αSMA expression, those with an intensity lower than 2500 a.u. as “low,” and those in between as “medium”.

### hiPSC-CF and hiPSC-CM Coculture

The effects of activated HCFs on cardiac functionality were assessed via the in direct coculture of HCFs and hiPSC-derived cardiomyocytes (hiPSC-CMs) in 6-well, flat bottom tissue culture plates (Genesee Scientific, 25-105). hiPSC-CMs were generated using a small molecule-based approach for inhibiting Gsk3 and Wnt, as described by Lian et al. in 2013.^27^ WTC11 hiPSCs, a GCaMP6f-expressing line kindly provided by Dr. Bruce Conklin, were differentiated into cardiomyocytes. This modification enables visualization of calcium transients during contraction via GFP fluorescence. GCaMP6f is a cpGFP-based sensor fused to calmodulin and the M13 peptide. Binding of Ca^2^+ to calmodulin’s EF-hand motifs induces a calmodulin–M13 interaction that stabilizes cpGFP in its fluorescent state, and dissociation upon Ca^2^+ removal returns it to low fluorescence. This reversible mechanism enables real-time monitoring of rapid Ca^2^+ dynamics during cardiomyocyte contraction. HCFs were seeded onto hydrogel disks printed at 70 mW/cm^2^, cultured in FGM-3 with 0.1 ng/mL TGFβ for 48 hours, and then placed into Millicell 6-well-hanging cell culture inserts with 1.0 μm pore size (MilliporeSigma, PLRP06H48). These inserts were then placed into wells containing hiPSC-CMs with the HCF-laden hydrogels submerged in the well’s medium without direct contact with the hiPSC-CM monolayer. Control conditions were generated by placing inserts with acellular 70 mW/cm^2^ gels in wells containing hiPSC-CMs. FGM-3 with 0.1 ng/mL TGFβ medium was used in all groups and exchanged every other day for 9 days. Assessment of calcium handling functionality of hiPSC-CMs based on GCaMP6f fluorescent signal was performed on days 0 (inserts added), 2, 5, and 9 of hiPSC-CM-HCF coculture. This analysis was done using a MATLAB computational pipeline we have previously reported on,^28,29^ wherein GFP signal intensity from time-stack images of hiPSC-CM contractions are used to evaluate hiPSC-CM excitation-contraction coupling metrics, specifically full-width half-max (FWHM), median absolute deviation of time-of-peak-arrival (MAD of TPA), and the decay constant.

### Statistical Analyses

All bar graphs depicted were generated using mean values with standard errors used to set the error bars. For comparisons with only two groups per analysis, a student’s t-test was used. A one-way ANOVA with pre-selected groups was chosen for comparisons with more than two groups.

## RESULTS

### Optimizing NorGel properties

Following the synthetic protocol by Van Hoorick et al. 2018,^25^ we successfully functionalized gelatin with norbornene to generate NorGel, as confirmed by the presence of the norbornene double bond peak (*δ* 6.22 – 5.90 ppm) in the ^1^H NMR spectrum of NorGel samples (Figure S2). Based on these ^1^H NMR spectra, we found an average of 42% amine functionalization with norbornene (0.163 mmol/g gelatin) (**Fig. 2A**, Equation S3). To determine the optimal crosslink density, we calculated the stoichiometric ratio of thiol groups (from 1,4-dithio-DL-threitol; DTT) to norbornene functionalities on NorGel using Equation S2. As an example, 0.2 g of NorGel (0.8 g PBS) required ∼2 mg of DTT (∼0.2 wt% of total mass) together with lithium phenyl(2,4,6-trimethylbenzoyl)phosphinate (LAP, 0.3 wt%) as photoinitiator and (2,2,6,6-tetramethylpiperidine-1-yl)oxidanyl (TEMPO, 0.1 wt% rel. to bulk) as stabilizer (**Fig. 2B**). Under a purely step-growth polymerization mechanism, a 1:1 thiol-to-ene molar ratio should result in the highest crosslink density. To confirm this, NorGel resins were prepared with varying amounts of DTT (0, 0.1, 0.2, 0.4, 0.6, 1, and 2 wt%) and examined using photorheology to determine the time to gelation upon light exposure (405 nm LED, 50 mW/cm^2^) and maximum storage modulus (*G’*) as a measure of stiffness (Figure S3-S4, Table S4). Increasing DTT from 0 to 0.2 wt% increased the final *G’* value, with 0.2 wt% being the highest. Specifically, no DTT (control) did not result in a change in *G’* (no reaction), while 0.1 and 0.2 wt% resulted in hydrogels with *G’* values of ∼0.3 ± <0.1 kPa and 19 ± 2 kPa, respectively, consistent with the stoichiometric functional group balance determined by ^1^H NMR. Increasing the concentration of DTT from 0.2 wt% to 2 wt% led to a reduction in the final modulus. Specifically, a concentration of 0.4 wt% and 2 wt% resulted in hydrogels with *G’* values of 12 ± 2 kPa and 1.8 ± 0.1 kPa, respectively. This decrease was attributed to an excess of thiol relative to ene functionalities, which generates DTT capped dangling chains, reducing the number of effective crosslinks and thereby lowering the overall crosslink density and stiffness. Next, we optimized the concentrations of LAP and TEMPO. We determined that 0.1 wt% TEMPO (rel. to bulk) was necessary to stabilize the sample and prevent gelation in the dark (Figures S5-S6, Table S5). Finally, a concentration of 0.3 wt% LAP (rel. to bulk) was selected as it achieved the fastest gelation (∼6.5 s) compared to 0.1 and 0.2 wt% LAP (Figures S7-S8, Table S6).

**Figure 2.**
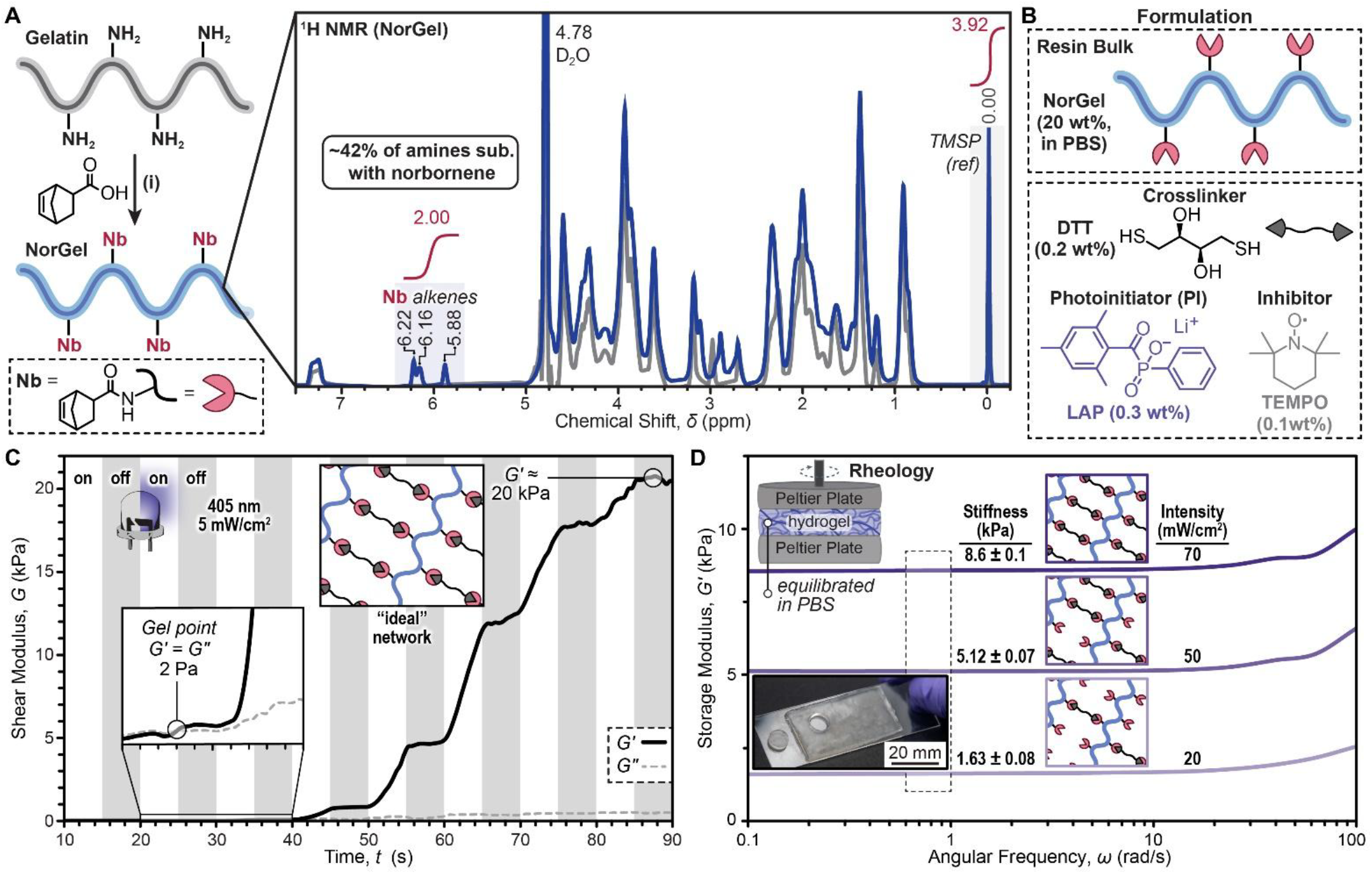
NorGel Synthesis and Hydrogel Network Characterization. A) Representative synthesis (left) and ^1^H NMR spectra (right) showing unmodified type B gelatin (gray) and the resulting norbornene-functionalized gelatin (blue). Highlighted areas represent regions used to quantify norbornene functionalization. (i) Conditions: EDC, NHS, DMSO, 20 hrs. B) Optimized resin formulation: 80 wt% PBS and 20 wt% NorGel bulk with 0.2 wt% DTT, 0.3 wt% LAP (photoinitiator), and 0.1 wt% TEMPO (stabilizer) to bulk. C) Temporal control over stiffness characterized using photorheology with 5 second light on-off cycling starting at 10 seconds. D) Stiffness of 3D printed norbornene-based hydrogels fabricated with different light dosages (20, 50, and 70 mW/cm^2^), equilibrated in PBS, and characterized using shear rheology. Inset image is of a 3D printed (50 mW/cm^2^) hydrogel with an 8 mm disk punched from it. *N* = 3 ± SD.

### Light intensity controls NorGel crosslinking kinetics and stiffness

Next, we assessed the effect of light intensity on hydrogel crosslinking kinetics via photorheology using a 405 nm LED at 20, 50, and 70 mW/cm^2^ intensities, with constant sample thickness of 100 µm and temperature of 37 °C. Increasing light intensity shortened the gel point, from 11.4 ± 1.3 s at 20 mW/cm^2^ to 6.5 ± 1.2 s at 50 mW/cm^2^ and 4.5 ± 1.2 s at 70 mW/cm^2^ (Figures S9-S12, Table S7). These results demonstrated that crosslinking kinetics could be modulated by adjusting the light intensity. We also investigated the effect of light dosage over time on *G’* using an on-off irradiation experiment (**Fig. 2C**). In this study, 20 wt% NorGel in PBS with optimized additives (0.3 wt% LAP, 0.1 wt% TEMPO, and 0.2 wt% DTT) held at 37 °C was exposed to violet light (405 nm, *I* = 5 mW/cm^2^) for 5-second intervals. For the first ∼20s after the initial light exposure, no change in *G’* was observed. However, at the fourth round of light exposure (∼300 mJ dosage), *G’* increased. Switching the light off resulted in a rapid *G’* plateau, indicating good temporal control, a prerequisite for grayscale 3D printing to tune stiffness.

Using a DLP 3D printer equipped with a 405 nm LED, rectangular samples (40 mm × 20 mm × 2 mm, *l* × *w* × *h*) were prepared with a constant layer thickness (100 µm) and exposure time (10 s) while varying light intensity (20, 50, and 70 mW/cm^2^). Following printing, the samples were equilibrated in PBS at 37 °C and punched into 8 mm disks for further rheological characterization and cell studies. Linear oscillatory shear rheology of the swollen hydrogels revealed that *G′* increased with light intensity, yielding 1.63 ± 0.08 kPa at 20 mW/cm^2^, 5.12 ± 0.07 kPa at 50 mW/cm^2^, and 8.60 ± 0.10 kPa at 70 mW/cm^2^ (**Fig. 2D**). These results demonstrated the ability to tune hydrogel stiffness by varying the dosage of light exposure during DLP 3D printing. Subsequent in vitro cell culture was performed using these three sample stiffnesses as described below.

### Efficient differentiation of HCF from hiPSCs

Following a published HCF differentiation protocol (**Fig. 3A**),^26^ we directed hiPSC differentiation towards an HCF fate. Brightfield imaging confirmed this transition, revealing hallmark morphological progression from pluripotent hiPSC colonies into spindle-shaped HCFs (**Fig. 3B**). HCF differentiation was confirmed at the protein level by immunofluorescence staining for the fibroblast marker vimentin. Concurrent alpha-smooth muscle actin (αSMA) staining identified the presence of activated HCFs. Day 20 differentiated cells exhibited qualitatively robust, nearly uniform vimentin expression, with a subset of cells exhibiting pronounced αSMA positivity, indicative of spontaneous activation (**Fig. 3C**, left). Further quantitative flow cytometry confirmed that over 96% of cells of the differentiated population expressed vimentin, demonstrating efficient hiPSCs differentiation into HCFs (**Fig. 3C**, right). Lastly, HCF differentiation was validated by gene expression analysis. qPCR analysis revealed statistically significant upregulation in the expression of *COL3A1*(20×10^8^-fold), *TCF21* (90-fold), and *TBX20* (8-fold) in day 20 HCFs relative to undifferentiated day 0 hiPSCs (**Fig. 3D, i-iii**). Increased *COL3A1* expression indicated significant upregulation of type III collagen production, suggesting that the cells in the differentiated population had increased ECM deposition. *TCF21* expression is essential for healthy cardiac fibroblast development^30^. Upregulation of this gene indicated that the differentiated population expressed CF-specific markers. Lastly, *TBX20* plays a critical role in heart development in several cell types within the cardiac niche, including cardiac fibroblasts^31,32^. Increased expression of *TBX20* further indicates that the day 20 differentiated cells were cardiac cells. Together, these results confirm the successful generation of a homogeneous HCF population.

**Figure 3.**
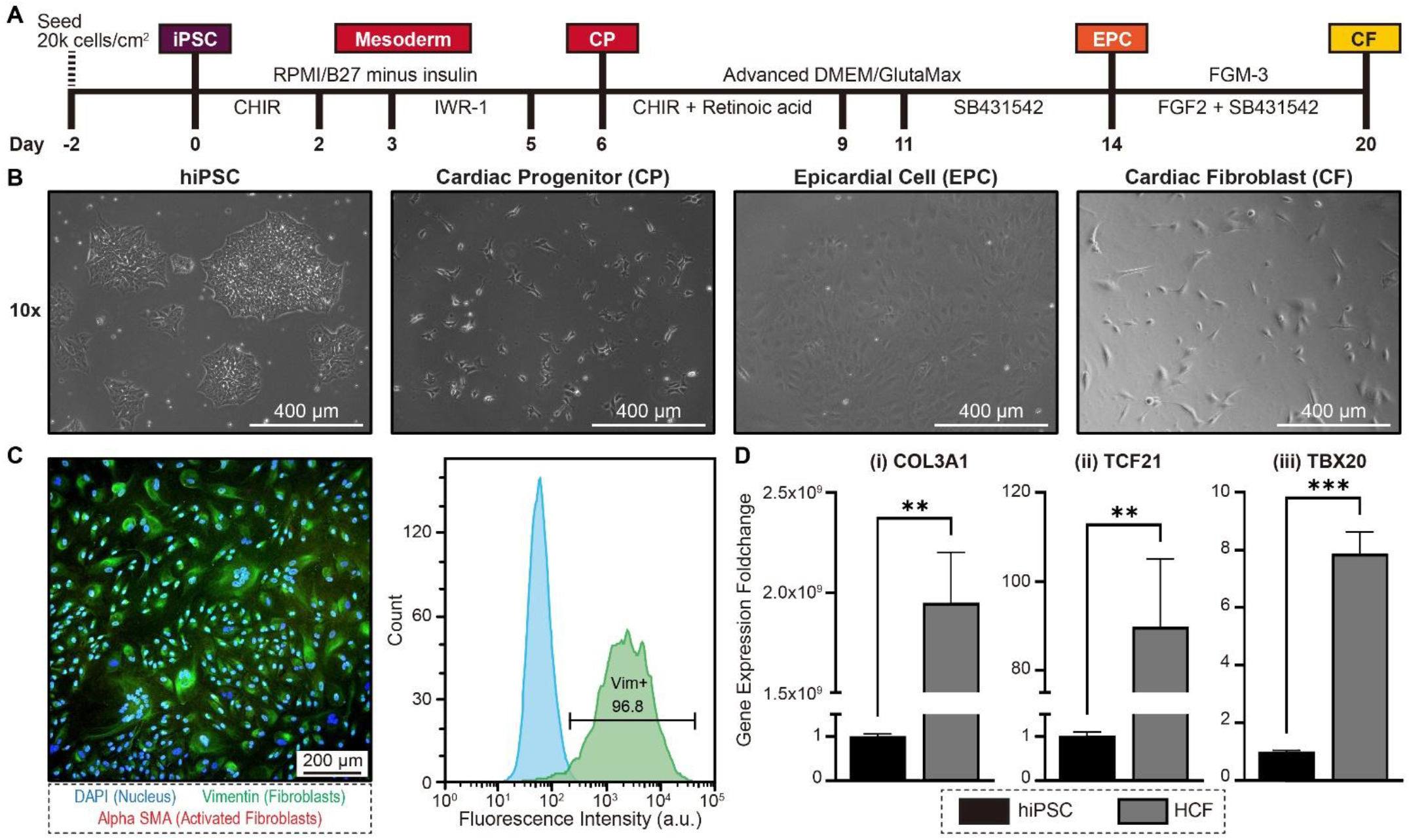
hiPSC-Cardiac Fibroblast Differentiation Protocol and Validation. A) The differentiation timeline for generating HCFs on day 20. B) Representative phase contrast images illustrating the transitions of changes in cell phenotypes during differentiation from hiPSCs to HCFs. C) day 20, hiPSC-derived HCFs displayed high differentiation efficiency and a quiescent phenotype, as indicated by positive vimentin staining and lack of α-smooth muscle actin (αSMA) expression, confirmed via microscopy (left) and flow cytometry (with hiPSCs used to set the negative gate). D) qPCR analysis of genes associated with a cardiac fibroblast phenotype in hiPSC-HCFs relative to hiPSCs, with expression foldchange calculated using the 2^-ΔΔCt^ approach). *N* = 3; ** and *** indicate p-values ≤ 0.01 and ≤ 0.001, respectively.

### HCFs transition from quiescent to activated state upon exposure to TGFβ on tissue culture plates

After establishing a source of HCFs, we confirmed that the derived cells exhibited a quiescent phenotype and could transition to an activated state upon exposure to TGFβ (**Fig. 4A**). The Quiescent state was assessed on HCFs cultured in fibroblast growth media (FGM-3) with and without SB431542 (a TGFβ inhibitor). To minimize activation from substrate stiffness, cells were plated on tissue culture plates coated with Geltrex. Flow cytometry revealed that there was no significant increase in the percent of αSMA-positive activated HCFs, whether SB431542 was present or not for 48 hours (**Fig. 4B**). Therefore, any increases in the abundance of activated HCFs following TGFβ exposure in subsequent experiments can be attributed to TGFβ promoting activation pathways. Notably, HCF exposure to either 0.1 ng/mL or 5 ng/mL TGFβ for 48 hours showed statistically significant increases in the abundance of activated HCFs relative to control samples not exposed to TGFβ. Although 5 ng/mL of TGFβ induced significantly greater HCF activation compared to lower dosages, 0.1 ng/mL of TGFβ was selected for HCF activation in subsequent studies using cell-laden hydrogel disks to avoid saturating the activation of HCFs and better elucidate the role that hydrogel stiffness plays in modulating HCF activation.

**Figure 4.**
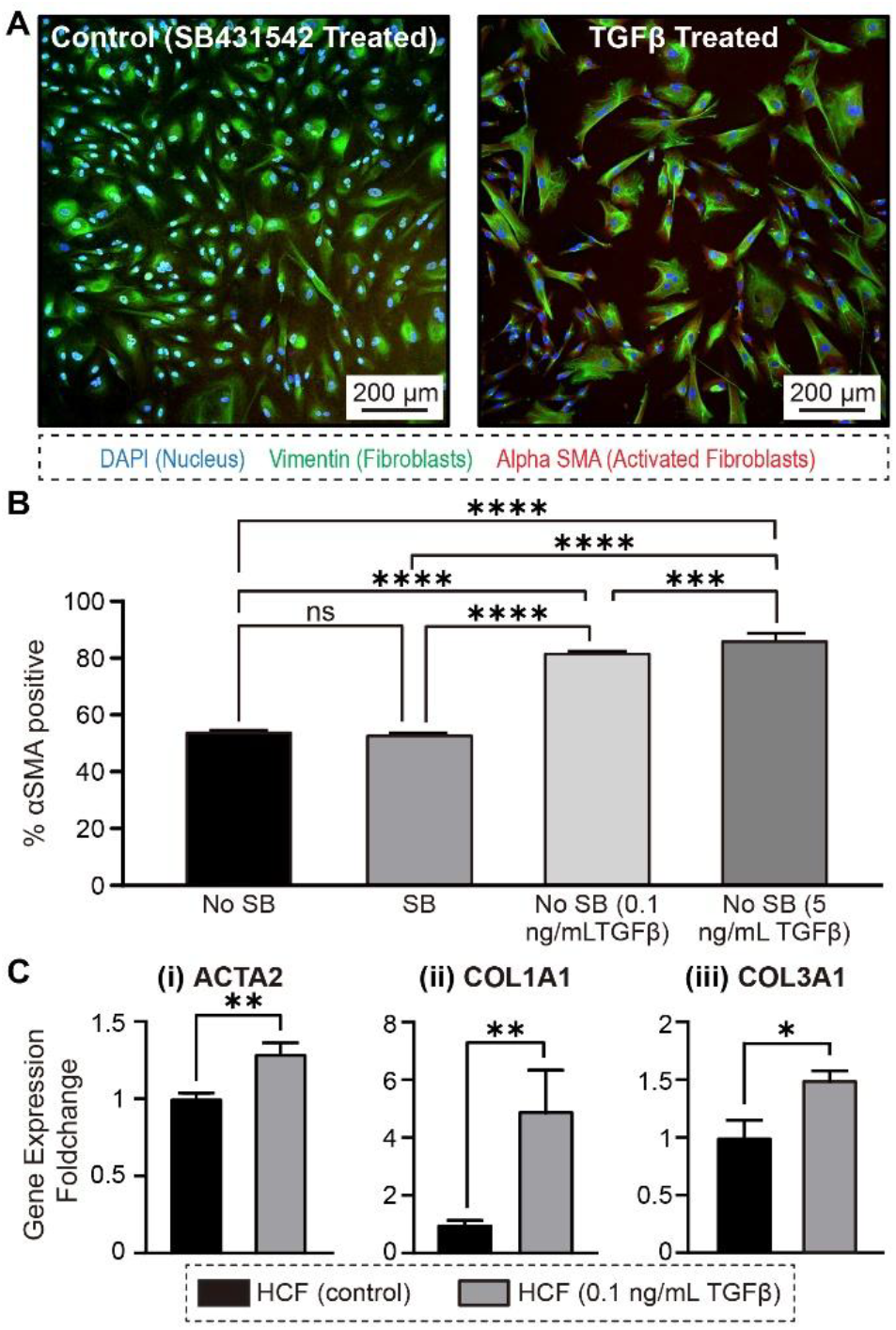
hiPSC-Cardiac Fibroblast Activation on Tissue Culture Plates. A) Day 20 HCFs were replated and cultured for 48 hours in FGM-3 supplemented with either SB431542 (left) or TGFβ (right), then stained for vimentin and αSMA. B) Flow cytometry data of the percent of activated fibroblasts (%αSMA+) cultured under four conditions: FGM-3 alone, FGM-3 with SB431542, FGM-3 without SB431542 with 0.1 ng/mL TGFβ, or FGM-3 without SB431542 with 5 ng/mL TGFβ. C) qPCR analysis comparing expression of genes associated with an activated cardiac fibroblast phenotype in HCFs treated with 0.1 ng/mL TGFβ compared to quiescent HCFs, with expression fold changes calculated using the 2^-ΔΔCt^ method and quiescent HCFs as the reference control population. Three wells were used for each condition (N=3); * and ** indicate p-values ≤ 0.05 and ≤ 0.01, respectively.

Next, we evaluated differences in the expression of *ACTA2, COL1A1*, and *COL3A1* genes in HCFs after 48 hours of exposure to 0.1 ng/mL TGFβ using qPCR analysis (**Fig. 4C**). A statistically significant 1.3-fold increase in the expression of *ACTA2*, a marker of fibroblast activation, was observed relative to the control, indicating that the TGFβ treatment promoted an activated phenotype in the HCF population. For *COL1A1* and *COL3A1* – genes responsible for producing collagen I and III, respectively – a 4.9 and 1.5-fold statistically significant increase in their expression was observed in the TGFβ treatment group, suggesting a transcriptional upregulation associated with ECM production in TGFβ activated HCFs. Overall, the observed protein and gene expression changes indicate that the differentiated HCFs respond to canonical fibroblast activation signaling and undergo activation.

### HCFs are activated by NorGel stiffness

After confirming that the differentiated HCFs could transition to an activated state, we next seeded the HCFs onto 3D printed hydrogel disks fabricated using light exposure intensities of 20, 50, or 70 mW/cm^2^, corresponding to *G’* values ranging from ∼1.5 – 8.5 kPa. Notably, this relative foldchange increase in stiffness is comparable to changes experienced by healthy cardiac tissue as it undergoes fibrosis, albeit with moderately lower absolute *G’* values. On day 20 of differentiation, HCFs were seeded onto the hydrogels of varying stiffnesses for 48 hours in FGM-3 to evaluate the impact of hydrogel stiffness on HSC activation. We also assessed the synergistic effect of coupling biophysical cues of substrate stiffness with biochemical stimulation of exposure to TGFβ (0.1 ng/mL) (**Fig. 5A**). As a control for the effect of TGFβ alone, we included HCFs exposed to TGFβ and cultured on thick Geltrex-coated tissue culture plates (*E* < 0.1 kPa)^33^, to prevent activation due to the underlying stiffness of standard tissue culture plates^34^. All groups were subsequently evaluated for changes in the expression of HCF activation-related genes (*ACTA2, COL1A1*, and *COL3A1*) (**Fig. 5B**). When examining differences in relative expression of *ACTA2*, all HCF populations cultured on hydrogels exhibited higher *ACTA2* expression compared to HCFs exposed to TGFβ and cultured on Geltrex coated plates, with a positive correlation between increased hydrogel stiffness and elevated *ACTA2* expression. Increased substrate stiffness alone (HCFs cultured on hydrogels without TGFβ) elevated *ACTA2* expression levels comparable to those observed in HCFs cultured on Geltrex-coated tissue culture plates and exposed to TGFβ, particularly for HCFs seeded on the 20 and 50 mW/cm^2^ hydrogels. Notably, the stiffest hydrogels (70 mW/cm^2^) induced a significantly greater increase in *ACTA2* expression. These results showed that our model can highlight the significant role substrate stiffness plays in activating cardiac fibroblasts. Moreover, the further activation of *ACTA2* expression in HCFs cultured on hydrogels and treated with TGFβ demonstrates that this model effectively recapitulates the synergistic effects of matrix stiffening and profibrotic signaling observed in cardiac fibrosis. Thus, the tunable stiffness of the 3D printed hydrogel models provides a robust platform for modeling the effects of stiffening cardiac scar tissue on cardiac fibroblasts, with stiffer hydrogels promoting greater expression of *ACTA2*.

**Figure 5.**
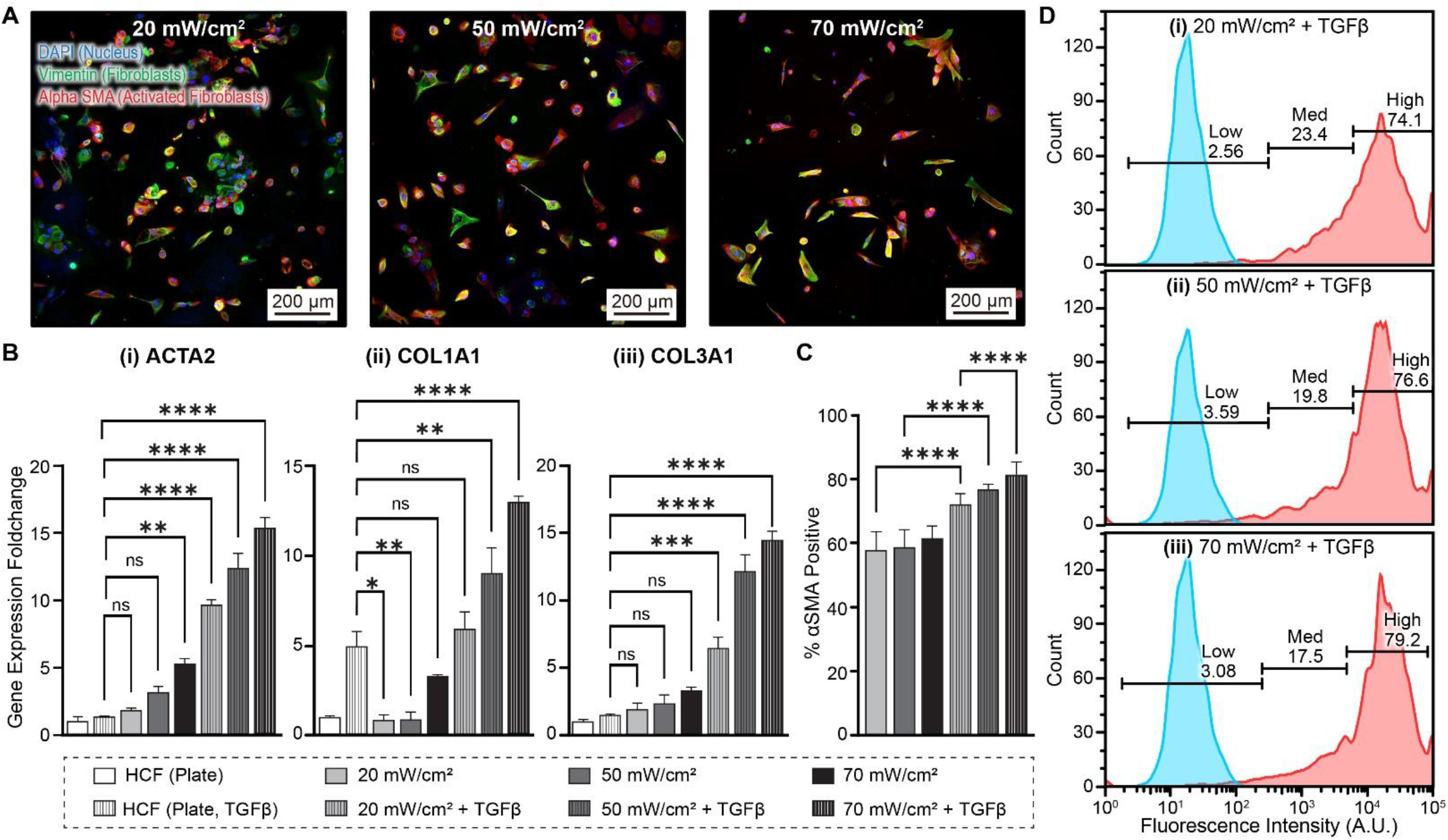
hiPSC-Cardiac Fibroblast Activation on 3D Printed Hydrogel Disks. A) Day 20 HCFs were replated onto 3D printed hydrogel disks fabricated using 20 mW/cm^2^ (left), 50 mW/cm^2^ (middle), or 70 mW/cm^2^ (right) light intensities, cultured for 48 hours in FGM-3 with 0.1 ng/mL TGFβ, and then stained for vimentin and αSMA. B) qPCR analysis comparing expression of genes associated with an activated cardiac fibroblast phenotype between quiescent HCFs (controls), HCFs cultured on tissue culture plates treated with TGFβ and HCFs cultured on hydrogels treated with or without TGFβ. Gene expression foldchange was calculated using the 2^-ΔΔCt^ method. C) Flow cytometry quantification of %αSMA+ in the same conditions as in panel B. D) Stratification of αSMA+ HCFs cultured on hydrogels and treated with TGFβ into “low” and “high” expression groups, based on fluorophore signal intensity, corresponding to lower and higher activation states, respectively. N=3 for each condition; *, **, ***, and **** indicate p-values ≤ 0.05, ≤ 0.01, ≤ 0.001, and ≤ 0.0001, respectively.

Hydrogel stiffness and *COL1A1* expression in HCFs showed a similar trend to that observed with *ACTA2* expression, where stiffer hydrogels promoted higher *COL1A1* expression (**Fig. 5B**). However, unlike *ACTA2*, increased hydrogel stiffness alone was not sufficient to significantly elevate *COL1A1* expression relative to TGFβ activation alone (HCFs cultured on Geltrex coated tissue culture plates). Specifically, TGFβ-activated HCFs on tissue culture plates demonstrated significantly higher *COL1A1* expression than HCFs cultured on the low-and intermediate-stiffness hydrogels (prepared with 20 & 50 mW/cm^2^ light) without TGFβ. In contrast, there was no significant difference in expression compared to HCFs cultured on the stiffest hydrogel (70 mW/cm^2^ light) without TGFβ. Among the TGFβ treated groups, HCFs cultured on low-stiffness hydrogels showed a non-significant increase in *COL1A1* expression, whereas those cultured on the intermediate and stiffest hydrogels exhibited statistically significant increases in *COL1A1* expression compared to TGFβ-treated HCFs cultured on Geltrex-coated tissue culture plates. Taken together, these data indicated that our cardiac fibrosis model successfully recapitulates key aspects of in vivo cardiac fibrosis, with increased *COL1A1* expression driven by both increased stiffness and the presence of inflammatory, profibrotic TGFβ. Notably, *COL1A1* expression did not fully parallel the trends observed with *ACTA2* expression, suggesting that while increased substrate stiffness alone can induce a significantly more activated HCF phenotype, substantial ECM gene expression is primarily driven by the presence of TGFβ. Similar trends were observed in *COL3A1*, with no significant differences in expression across the TGFβ-free hydrogel conditions and significant increases in all TGFβ treated hydrogel conditions compared to HCFs cultured and activated on Geltrex-coated tissue culture plates. Accordingly, these data suggest that our cardiac fibrosis model of HCFs cultured on 3D printed hydrogels can recapitulate transcriptomic changes in HCFs that mimic the activation of cardiac fibroblasts during cardiac fibrosis.

Further analysis of HCF activation at the protein level using flow cytometry after αSMA staining revealed that TGFβ exposure led to significant increases in the abundance of activated HCFs across all hydrogel stiffness conditions (**Fig. 5C**). Furthermore, quantifying the fluorescent intensity from the fluorophore-conjugated αSMA antibody showed a positive correlation between hydrogel stiffness and the percent of HCFs exhibiting “high” αSMA expression (**Fig. 5D**). These findings confirm that both TGFβ and increased substrate stiffness contribute to HCF activation at the protein level, complementing the transcriptomic data.

### HCF activation contributes to hiPSC-CM dysfunction

Next, we cocultured activated HCFs on the stiffest 3D printed hydrogels with hiPSC-CMs, which were separately differentiated in a monolayer on tissue culture plates, to assess whether our HCF models could mimic the pathologic effects of cardiac fibrosis (**Fig. 6A**). Notably, the HCF-laden hydrogels were not in direct contact with the differentiated hiPSC-CMs, ensuring that all interactions occurred via soluble factors in the shared media. The functionality of hiPSC-CMs was evaluated using excitation-contraction coupling metrics that were quantified based on calcium ion transient-actuated fluorescence (**Fig. 6B** and Movie S1). Key metrics extracted include the duration, decay constant, and synchronicity of hiPSC-CM contractions. The duration of hiPSC-CM contractions was measured by the calcium transient signal’s full-width half-max (FWHM), where a smaller FWHM indicates a more functional and mature myocyte. Both the cocultured hiPSC-CMs and hiPSC-CMs with acellular hydrogels (control) displayed decreased FWHM values over the course of the experiment. However, the cocultured group showed no significant differences in FWHM values compared to controls on any day group except for a transient change on day 5, which was not sustained through day 9 (**Fig. 6C, i**), indicating that hiPSC-CM contraction duration was not disrupted by coculture with activated HCFs in the long term. As a second metric, decay constants, which provide a measure of how rapidly myocytes relax before contracting again, were quantified, with lower values indicating faster relaxation and, consequently, more functionally mature cardiomyocytes (**Fig. 6C, ii**). Similar to duration, there was no significant difference in decay constant between control hiPSC-CMs and those exposed to the cardiac fibrosis model by the end of the 9 days of coculture. Notably, differences emerged in the last excitation-contraction coupling metric analyzed, the synchronicity of contractions. Synchronicity was quantified by the median absolute deviation of time-of-peak-arrival (MAD of TPA) of calcium transient signals, with a lower MAD of TPA indicating greater synchronicity and, in turn, more functionally competent cardiac tissue (**Fig. 6C, iii**). Control hiPSC-CMs demonstrated a decrease in MAD of TPA over the 9 days of culture, indicating that additional time in culture allowed for improved myocyte synchronicity. In contrast, the hiPSC-CMs cocultured with activated HCFs displayed a reduction in MAD of TPA from day 0 to day 2 relative to the control hiPSC-CMs. Though this difference was not statistically significant, it does suggest that coculture with activated HCFs may have temporarily improved synchronicity in hiPSC-CMs. However, this improvement in synchronicity was short-lived, as MAD of TPA of the hiPSC-CMs exposed to activated HCFs on hydrogels proceeded to display increased MAD of TPA from day 2 to day 9, with day 9 hiPSC-CMs in the treated group having a statistically significant increase in MAD of TPA compared to day 9 hiPSC-CMs in the control group. This transient improvement in synchronicity, followed by a significant decline, may be attributed to cytokines secreted by activated HCFs. The activated cardiac fibroblast secretome can impact cardiomyocyte behavior, such as inducing cardiomyocyte hypertrophy.^35,36^ Changes in hiPSC-CM behavior, like cell hypertrophy, may have had an initial beneficial effect on synchronicity. However, extended coculture and prolonged exposure to the activated HCF secretome could have recapitulated the pro-inflammatory environment following cardiac injury, ultimately resulting in a deleterious effect on myocyte synchronicity.

**Figure 6:**
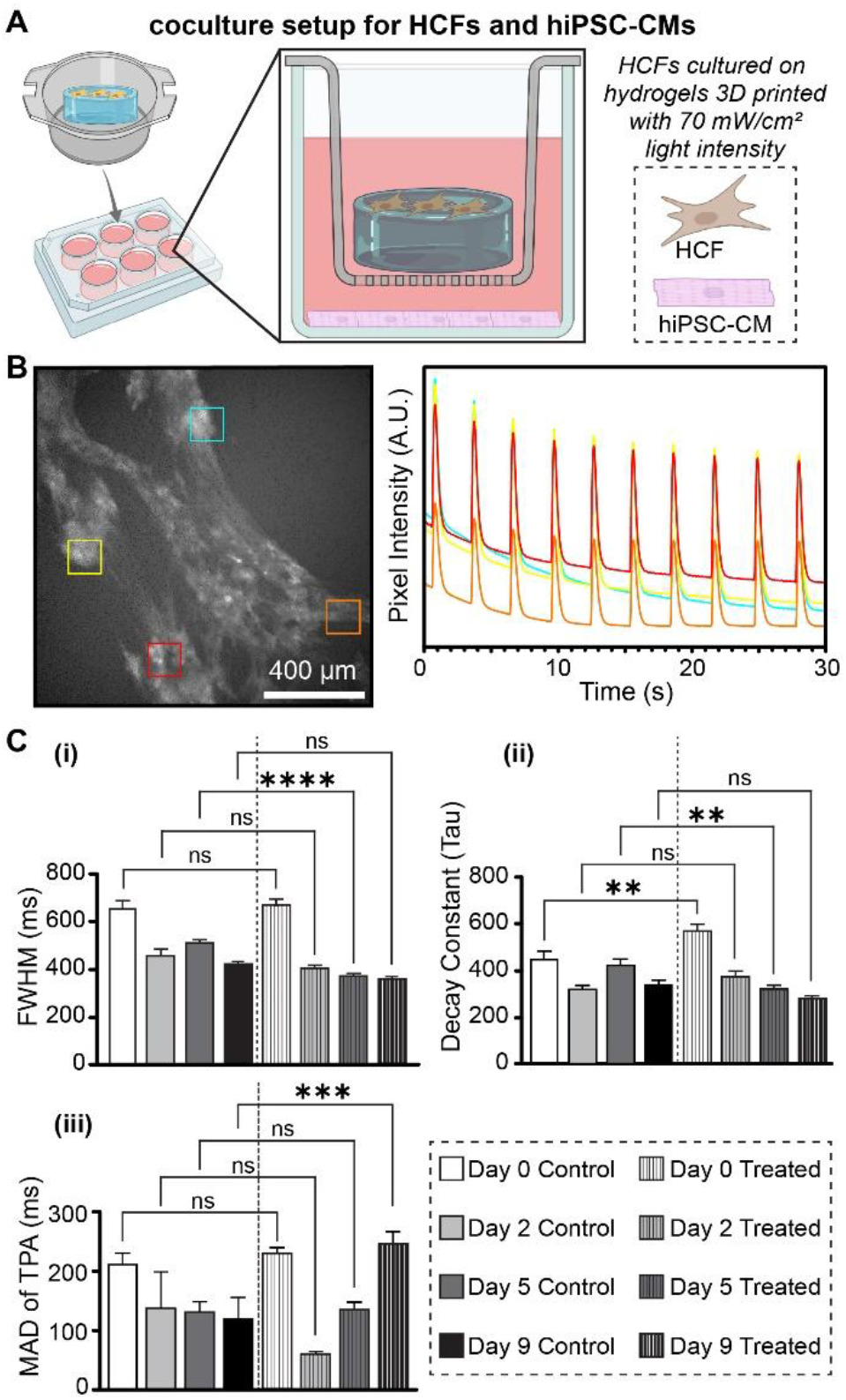
Effects of Activated hiPSC-cardiac Fibroblasts on hiPSC-cardiomyocyte Function. A) Graphical representation of the coculture setup for HCFs and hiPSC-CMs. HCFs were cultured on hydrogels crosslinked with 70 mW/cm^2^ light intensities (made with BioRender.com). B) Time-stack fluorescence images analyzed using a MATLAB-based computational pipeline. Regions of interest (ROIs) were selected (left) to extract calcium traces (right), enabling quantification of electromechanical coupling in hiPSC-CMs via calcium transient-actuated GCaMP fluorescence signal. C) Electrochemical coupling metrics extracted from the MATLAB pipeline: (i) full-width half-max (FWHM; a measure of contraction duration), (ii) decay constant (a measure of calcium ion reuptake), and (iii) median absolute deviation of time-of-peak-arrival (MAD of TPA; a measure of contraction synchronicity). 3 ROIs were taken for each hydrogel-seeded disk, and a total of 3 hydrogel-seeded disks were tested for each condition. **, ***, and **** indicate p-values ≤ 0.01, ≤ 0.001, and ≤ 0.0001, respectively.

## DISCUSSION

The heart’s limited regenerative capacity is one of the most significant barriers to addressing cardiovascular disease. Overcoming this challenge requires a deeper understanding of natural cardiac wound healing processes to identify the heart’s inherent limitations and develop strategies to compensate for them. Specifically, it is essential to investigate cardiac fibrosis, a condition that emerges when native repair processes shift from being beneficial to pathological due to inadequate regulation of cardiac fibroblast-mediated repair pathways. Therefore, it is critical to investigate cardiac fibroblasts and their behavior in the environment of a cardiac injury, as cardiac fibroblasts activated by signaling from injured and scarred cardiac tissue are responsible for the onset of cardiac fibrosis and its progression toward heart failure. Such investigations can significantly benefit from in vitro models that recapitulate the native environment of interest.

Previously developed models of cardiac fibrosis have either examined only biochemical stimuli of cardiac fibrosis,^37^ utilized primary cardiac fibroblasts,^38,39^ or manipulated mechanical properties outside of cell substrate stiffness.^39,40^ While these models are valuable, there are gaps that they do not address. Simultaneous biochemical and mechanical stimuli play a key role in cardiac fibroblast activation in cardiac fibrosis,^15,22^ making it critical to include both means of stimulus when developing an in vitro model of cardiac fibrosis. Differentiating cardiac fibroblasts from hiPSCs rather than sourcing them from primary cell lines is beneficial for scalability and potential personalized medicine applications. Lastly, discrete tuning of cell substrate stiffness is vital to unravel the explicit effects of increased stiffness on cardiac fibroblast activation, as ECM stiffening is a key fibroblast activator in cardiac fibrosis.^8,15^ Accordingly, we present our efforts to develop an in vitro model of cardiac fibrosis using 3D printed hydrogels of tunable stiffness with cardiac fibroblasts derived from hiPSCs with both discrete and simultaneous biochemical and mechanical stimuli to induce fibroblast activation.

We have successfully synthesized norbornene functionalized gelatin and crosslinked it with DTT via a temporally and dose-controlled photopolymerization using LAP as the photoinitiator excited by 405 nm light in 80 wt% PBS (**Fig. 2A, 2B**). By varying the intensity of irradiated light, we tailored the crosslinking density and, in turn, the stiffness of the resulting hydrogel (**Fig. 2D**). Using different intensities of 405 nm light, we created 3D printed hydrogel disks with a range of stiffness (1.63 ± 0.08 kPa (20 mW/cm^2^), 5.12 ± 0.07 kPa (50 mW/cm^2^), and 8.59 ± 0.13 kPa (70 mW/cm^2^), matching the foldchange increase in stiffness between healthy and fibrotic cardiac tissue (**Fig. 2D**). The tunable stiffness of the 3D printed hydrogels was a key component in developing our cardiac fibrosis model, as substrate stiffness plays a significant role in activating cardiac fibroblasts and promoting cardiac fibrosis.

A second key aspect of our cardiac fibrosis model involved generating a reliable source of human cardiac fibroblasts. To that end, we utilized HCFs derived from hiPSCs, as hiPSCs offer a more proliferative and renewable cell source than commercially available primary human cardiac fibroblasts while preserving donor-specific traits essential for in vitro models. We were able to generate quiescent HCFs via directed differentiation from hiPSCs and subsequently activate said HCFs both in tissue culture plates and on 3D-printed hydrogel disks. Comparing the activation of HCFs on the two different culture substrates revealed significant differences in the increase in fibroblast activation markers, with stiffer hydrogel disks promoting significantly greater activation, as evidenced by both gene expression and flow cytometry analyses. Because the convergence of TGFβ signaling and increased substrate stiffness is critical for driving cardiac fibroblast activation to pathological levels, our 3D-printed hydrogel model’s ability to recapitulate this important interplay highlights its utility as a platform to study cardiac fibroblast activation. In addition, we demonstrated that our model can replicate the impact of native cardiac fibrosis on cardiac function, as evidenced by changes in hiPSC-CM excitation-contraction coupling metrics. By analyzing calcium traces with our computational pipeline, we revealed that hiPSC-CM synchronicity was significantly impaired in cocultures of activated HCFs with hiPSC-CMs when compared to hiPSC-CMs cultured acellular hydrogels as controls. Rhythmic, synchronized beating of CMs is essential for healthy cardiac output. As such, the decreased synchronicity observed in our coculture condition indicates that our cardiac fibrosis model caused a loss of cardiac function in the monolayer of hiPSC-CMs. Furthermore, while the other excitation-contraction coupling metrics that were analyzed did not display significant differences between control and treated hiPSC-CMs, it is important to note that the coculture approach used did not allow for direct cell-cell contact between HCFs and hiPSC-CMs. This may explain why more pronounced reductions in analyzed metrics were not observed, as cell-cell contact would enable activated HCFs to deposit ECM proteins that directly interface with hiPSC-CMs, potentially affecting their behavior. Moving forward, a more comprehensive model of cardiac fibrosis should incorporate coculture, where both myocytes and activated fibroblasts are localized on (or within) the same substrate to better mimic the fibrotic microenvironment.

## CONCLUSION

Herein, we report the development of cardiac fibroblast tissue constructs fabricated using hiPSC-derived cardiac fibroblasts and 3D-printed hydrogels of tunable stiffness. This tunability enabled HCFs to be cultured on substrates of increasing stiffness to examine the impact of substrate stiffness on cardiac fibroblast activation. Stiffer substrates resulted in an increase in the abundance of activated HCFs in the population as well as an increase in the expression of markers of fibroblast activation, mimicking the behavior of native cardiac fibroblasts undergoing activation in an injured heart. Furthermore, by combining substrates of varying stiffness with a constant dose of TGFβ, our model demonstrates that it is important to recapitulate the conjunction of biochemical and biomechanical cues in native cardiac fibrosis to induce a significant shift towards an activated phenotype. Lastly, we demonstrated that the shift in HCF activation induced by our model was sufficient to produce measurable effects on the functionality of hiPSC-derived cardiomyocytes. These findings suggest that our constructs effectively recapitulate both quiescent and activated cardiac fibroblast behavior, with the potential for further development to more accurately model activated fibroblast dynamics in cardiac fibrosis and their effects on cardiac function.

## Supporting information

SupportingInformation

MovieS1_BeatingCM

## ACKNOWLEDGEMENTS

The authors acknowledge and appreciate support from the following funding sources: Harry S. Moss Heart Trust (UTAUS-FA00002510; S.S., N.M., L.M.S., J.H., Y.L., M.T.K., Z.A.P., and J.Z.), National Heart, Lung, and Blood Institute F31 Grant (5F31HL170717-03; S.S.), NSF Graduate Research Fellowship Program (DGE-2137420; E.A.R.), and partial support from the Robert A. Welch Foundation (F-2007; Z.A.P.). The authors acknowledge Marcel Lopez Reed, Sneha Sinha, and Maanas Gupta for their support with experimental efforts.

