## SupportingInformation for "Modulating Hydrogel Stiffness Through Light-Based 3D Printing to Mimic Cardiac Fibrosis and Cardiomyocyte Dysfunction Using hiPSC-Derived Cells"

**This PDF file includes:**

Methods
Supplementary Text
Figures S1 to S12
Tables S1 to S7

**Table of Contents**

**S1. Instrumentation** ........ **S3**

***S1.1. Supplemental Methods*** ........ ***S3***

***S1.2. NorGel Synthesis*** ........ ***S4***

**S2. Supplementary Text** ........ **S6**

***S2.1. Photorheology*** ........ ***S6***

***S2.1.1. Determination of optimal DTT content (wt%)*** ........ ***S6***

***S2.1.2. Determination of optimal TEMPO content (wt%)*** ........ ***S7***

***S2.1.3. Determination of optimal LAP content (wt%)*** ........ ***S8***

***S2.1.4. Light intensity effect on gel point*** ........ ***S9***

**S3. Supplementary Tables** ........ **S11**

**Reference** ........ **S13**

### S1. Instrumentation

#### *S1.1. Supplemental Methods*

*Nuclear Magnetic Resonance (NMR) Spectroscopy*. NMR Spectra were recorded on an Agilent MR 400 MHz spectrometer utilizing $D_2O$ with 0.05wt% TMSP as the solvent. $^1H$ NMR were carried out coupled and referenced to the $D_2O$ chemical shift at 4.79 ppm.

*Lyophilizer*. A Labconco FreeZone 2.5 Liter -84 °C Benchtop Freeze Dryer (Cat. Number 710201000) equipped with a combination vacuum pump (**#:** 7584002) from Labconco. The typical pressure was <3 x $10^{-3}$ mBar and the typical temperature was -80 °C to remove water from NorGel and GelMA samples after dialysis. Gelatin-based samples suspended in water were poured into glass containers (Labconco Fast-Freeze™ 7541100 Clear Glass Drying Flask, 750 mL) and capped with rubber lids (#7541100) equipped with ¾" diameter holes for use with ¾" diameter glass rods to attach to the instrument. The typical time to remove water was 72 hours.

*Solvent Purification System*. Vacuum Atmospheres Solvent Purifier system, VAC 103991 removes water by circulating solvent using a pump through a solid-phase desiccant (Part number: 105643, 105644, 105646, 105649, 105652, 105655, and105653) housed in stainless-steel cartridges.

*DLP 3D Printer.* 3D printing was performed using a custom-made, digital light processing (DLP) 3D printer (MONO3MZ2, Monoprinter). The printer contains an LED projector with a wavelength centered at 405 nm (PDC04-405 nm). The projector resolution is 1920 × 1080 pixels, with each pixel being 23.2 μm × 23.2 μm at the image plane. Digital files for each LED projection were generated by designing the 3D models using computer-aided design (CAD) software (Blender, SolidWorks) and exported into an STL format. Each STL file was imported into MonoWare software. Then, the selected 3D model was sliced into multiple 2D MNF files with a slice thickness of 100 μm. Each image from the MNF files were then projected onto the bottom of a resin vat with maximum image plane dimensions of 25 mm × 44 mm. A transparent fluorinated ethylene propylene (FEP) polymer film (Teflon FEP film, 127 μm thick, DuPont) was used as the base of the resin vat to provide a non-stick bottom surface. The resin vat was equipped with an ITO coated glass bottom and a metal holder that was heated to 37°C.

*Light Sources.* For all experiments aside from 3D printing, the following LED was used: a violet Type B LED (cat. no. LCS-0405-12-22, Mightex Systems) with an emission centered at ~405 nm and a full width at half maximum (FWHM) of 14.0. This LED was used in combination with a current-adjustable driver (SLC-MA02-U, Mightex Systems) for intensity control, such that intensity between experiments could be matched. Irradiation intensity was measured with a ThorLabs PM100D photometer equipped with either a silicon-based photodiode power sensor (S120VC and S130C) or thermal power sensor (S401C and S175C) depending on the intensity of the light used.

### *S1.2. NorGel Synthesis*

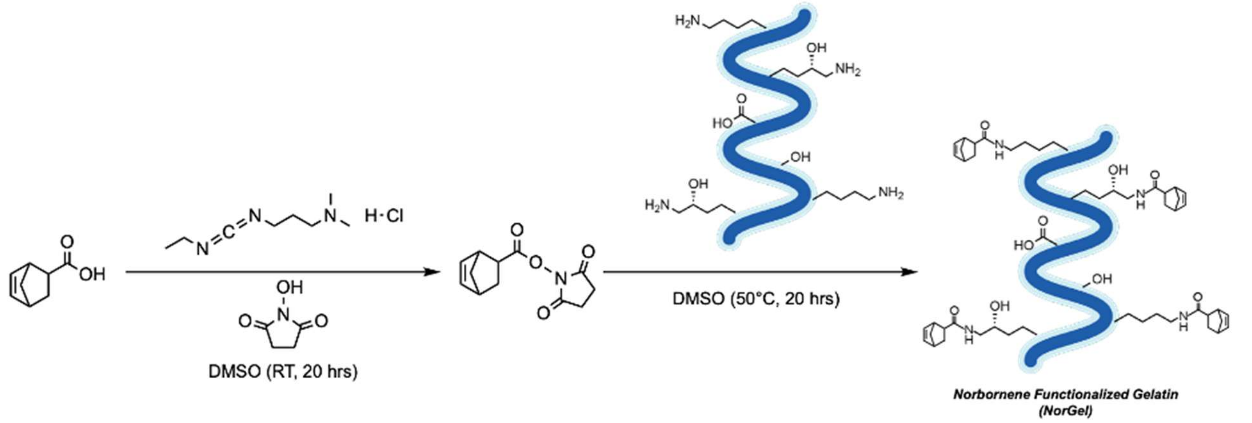


**Figure S1. A schematic representing the** synthetic pathway followed in this study to functionalize Type B Gelatin with norbornene carboxylic acid.

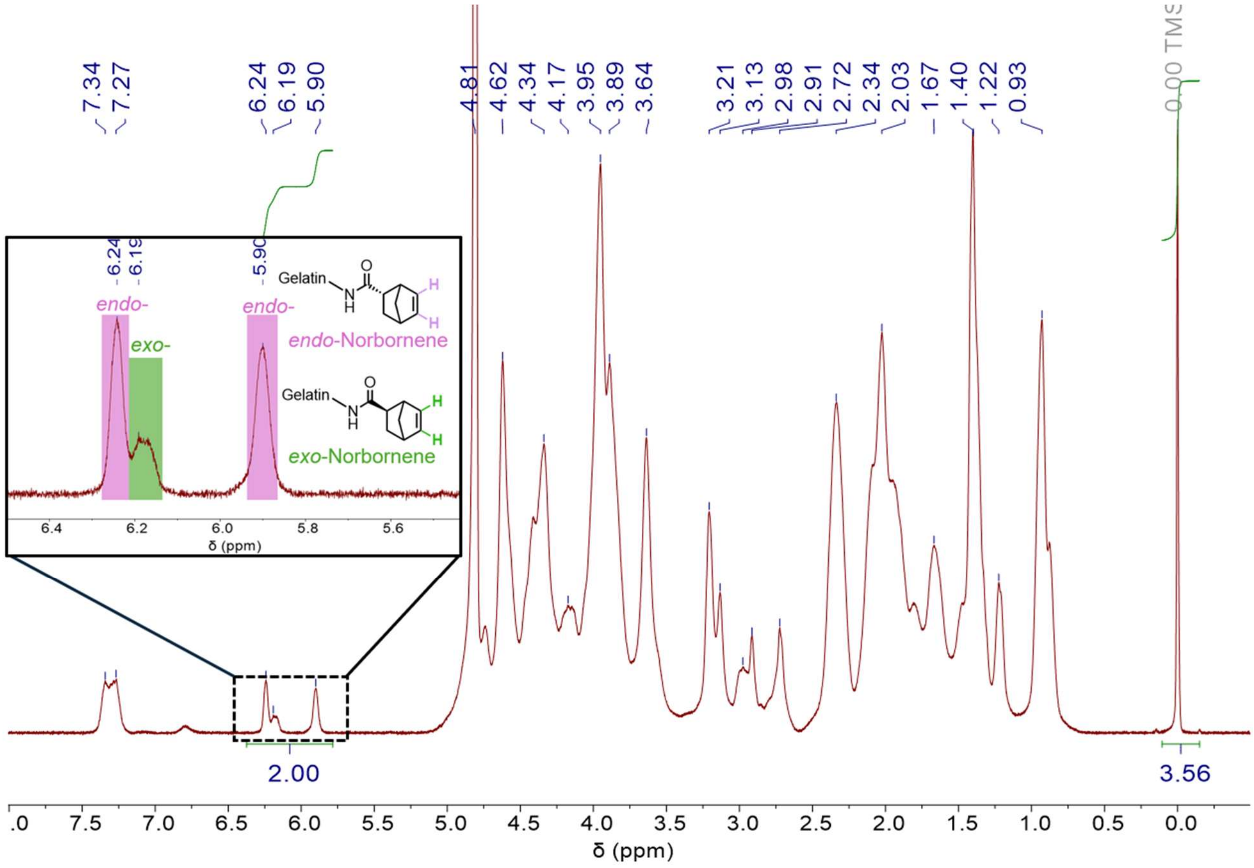


**Figure S2.** Representative $^{1}$H NMR spectrum of modified norbornene-functionalized Type b gelatin (NorGel) in $D_2O$ (referenced at 4.79 ppm) with 0.05 wt% TMSP. The norbornene peak at 6.24-5.90 ppm was integrated and set to 2 as the reference and the TMSP peak at 0.00 ppm was integrated.

The moles of norbornene (Nb) per grams of NorGel was determined using the following equations:

$$\frac{mol_{Nb}}{m_{NorGel}} = \frac{mol_{TMSP} \times \frac{H_{TMSP\ (0\ ppm)}}{I_{TMSP\ (0\ ppm)}} \times \frac{H_{Nb\ (6.24-5.9\ ppm)}}{I_{Nb\ (6.24-5.9\ ppm)}}}{m_{NorGel}}$$ **Equation S1**

where mol is moles, *m* is mass (in grams), *H* is number of protons (9 for TMSP, 2 for Nb), and *I* is integration (set to 2 for Nb).

Following the calculation of moles of Nb per grams of NorGel, the amount of dithiothreitol (DTT, *f* = 2) needed for "ideal" step growth polymerization could be determined using the following equation:

$$m_{DTT} = m_{NorGel} \times \frac{mol_{Nb}}{m_{NorGel}} \times \frac{0.5\ mol_{DTT}}{1\ mol_{Nb}} \times MW_{DTT}$$ **Equation S2**

where mol is moles, *m* is mass (in grams), and *MW* is molecular weight (in g/mol).
For each batch of Nb-functionalized gelatin, the degree of functionalization was calculated by dividing the measured Nb content (mmol/g gelatin) by the theoretical amine density of gelatin (0.385 mmol primary amines/g gelatin),[S1] based on the assumption that the EDC/NHS coupling occurs between 5-norbornene-2-carboxylic acid and the primary amines of gelatin, as has been reported previously.[S2]

$$Degree\ of\ Functionality\ (\%) = \frac{Nb\ (mmol/\ g\ gelatin)}{0.385\ (mmol/\ g\ gelatin)} \times 100\%$$ **Equation S3**

### S2. Supplementary Text

#### *S2.1. Photorheology*

All samples were measured at 37 °C.

##### *S2.1.1. Determination of optimal DTT content (wt%)*

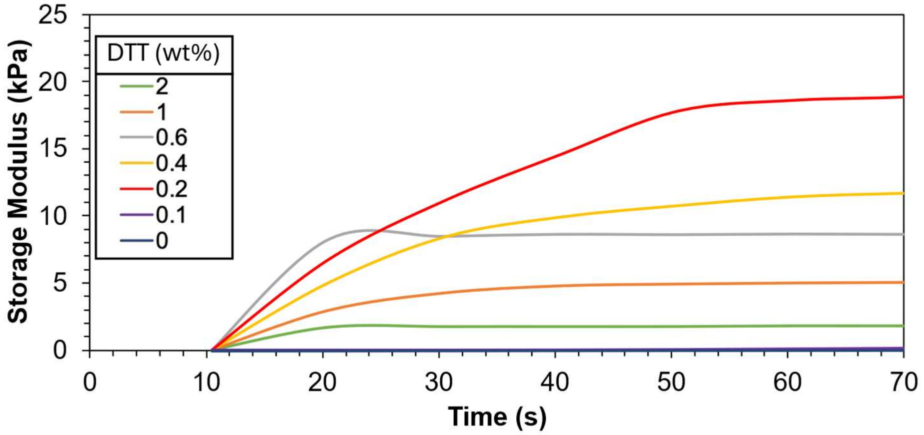


**Figure S3.** NorGel photorheology data showing representative storage modulus vs. time plot as a function of DTT concentration (provided in wt% relative to bulk). Samples consisted of 20 wt% NorGel in 80 wt% PBS, 0.1 wt% LAP and 0.1 wt% TEMPO and irradiating with a violet LED (405 nm LED, 50 mW/cm2, turned on at 10 s). Sample thickness was 100 µm and samples were sealed with mineral oil to prevent evaporation. Notably, the wt% of DTT determined to be optimal (0.2wt%) corresponded to the amount calculated for ideal stoichiometric conversion from $^1$H NMR spectroscopy. $N = 3 \pm$ SD.

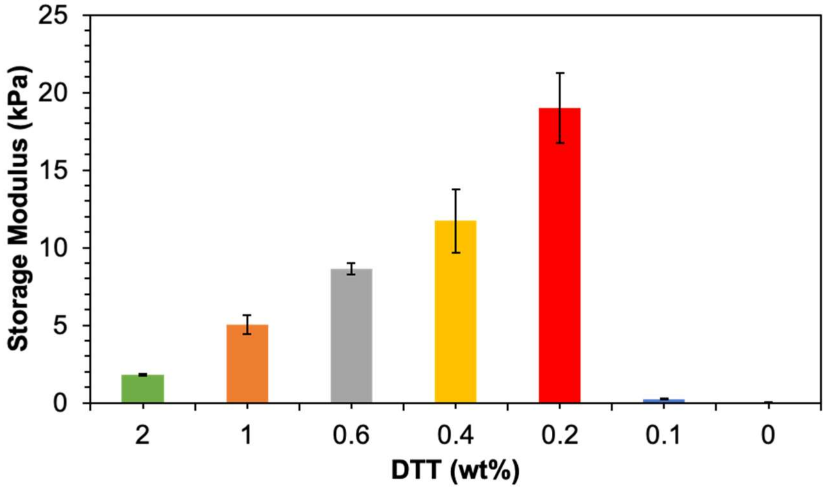


**Figure S4.** Compiled data for NorGel storage modulus values (at 60s light exposure) as a function of DTT concentration (wt% relative to bulk). Samples consisted of 20 wt% NorGel in 80 wt% PBS, 0.1 wt% LAP and 0.1 wt% TEMPO, and irradiated with a violet LED (405 nm LED, 50 mW/cm2). Sample thickness was 100 µm and samples were sealed with mineral oil to prevent evaporation. Notably, the wt% of DTT determined to be optimal (0.2wt%) corresponded to the amount calculated for ideal stoichiometric conversion from $^1$H NMR spectroscopy. $N = 3 \pm$ SD.

***S2.1.2. Determination of optimal TEMPO content (wt%)***

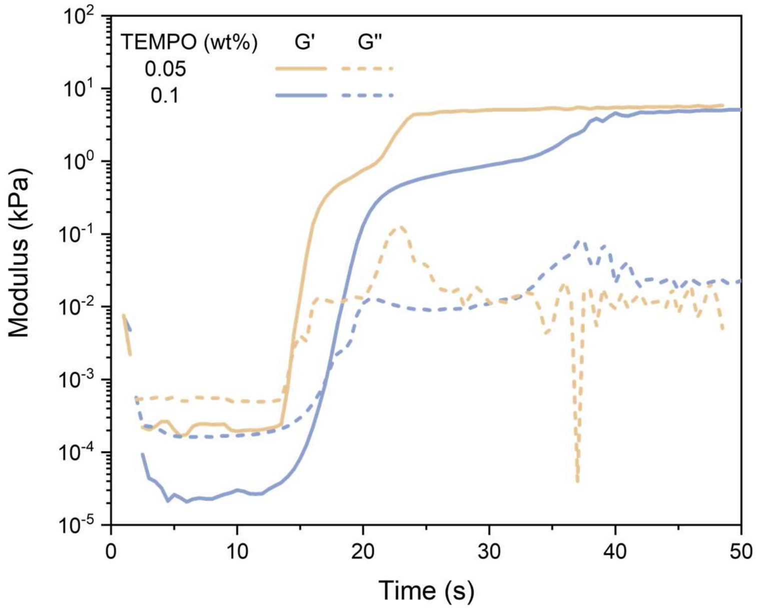


**Figure S5.** NorGel photorheology data showing representative storage modulus vs. time plot as a function of TEMPO concentration (provided in wt% relative to bulk). Samples consisted of 20 wt% NorGel in 80 wt% PBS, 0.2 wt% DTT and 0.3 wt% LAP, and irradiating with a violet LED (405 nm LED, 50 mW/cm$^2$, turned on at 10 s). Sample thickness was 100 µm and samples were sealed with mineral oil to prevent evaporation. added. Above 0.1wt% TEMPO, inhibition times >30s were observed and therefore not considered for future testing. Notably, 0.05 wt% TEMPO samples were not stable, showing curing in the absence of light exposure within 1 hour of mixing, precluding use in 3D printing. As a result, concentrations below 0.05 wt% TEMPO were not considered.

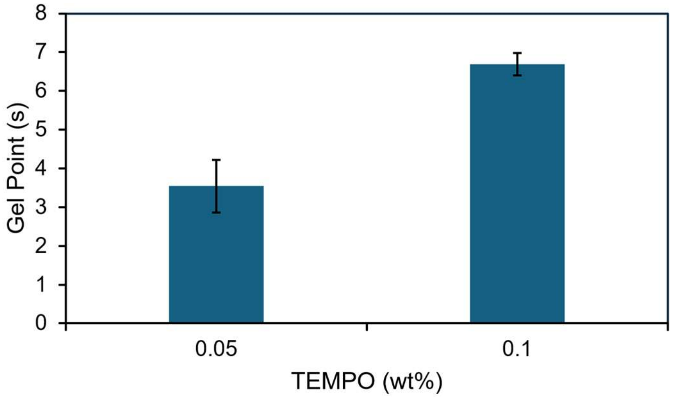


**Figure S6.** Effect of TEMPO concertation on NorGel point of gelation Samples consisted of 20 wt% NorGel in 80 wt% PBS, 0.2 wt% DTT and 0.3 wt% LAP, and irradiated with a violet LED (405 nm LED, 50 mW/cm$^2$). Sample thickness was 100 µm and samples were sealed with mineral oil to prevent evaporation. $N = 3 \pm$ SD.

***S2.1.3. Determination of optimal LAP content (wt%)***

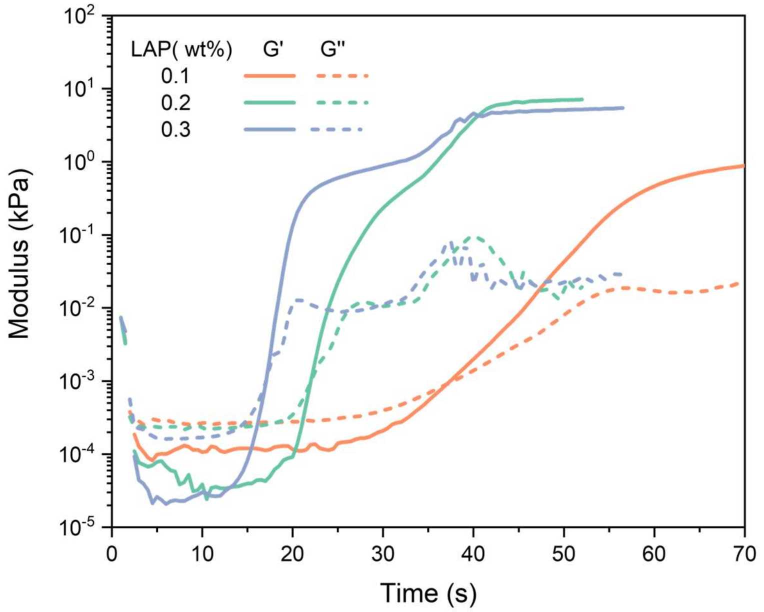


**Figure S7.** NorGel photorheology data showing representative storage modulus vs. time plot as a function of LAP concentration (provided in wt% relative to bulk). Samples consisted of 20 wt% NorGel in 80 wt% PBS, 0.2 wt% DTT and 0.1 wt% TEMPO, and irradiating with a violet LED (405 nm LED, 50 mW/cm$^2$, turned on at 10 s). Sample thickness was 100 µm and samples were sealed with mineral oil to prevent evaporation.

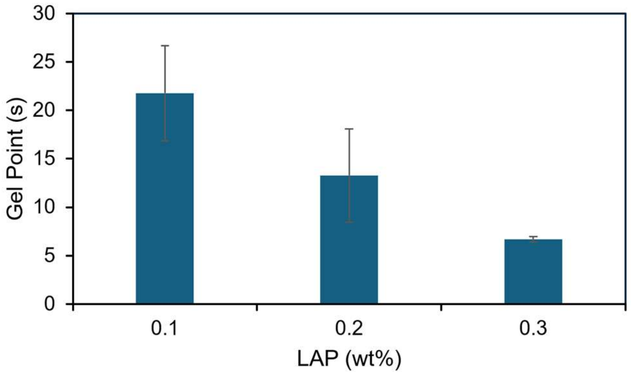


**Figure S8.** Effect of LAP concentration (wt% relative to bulk) on NorGell point of gelation. Samples consisted of 20 wt% NorGel in 80 wt% PBS, 0.2 wt% DTT and 0.1 wt% TEMPO, and irradiated with a violet LED (405 nm LED, 50 mW/cm$^2$). Sample thickness was 100 µm and samples were sealed with mineral oil to prevent evaporation. $N = 3 \pm$ SD.

***S2.1.4. Light intensity effect on gel point***

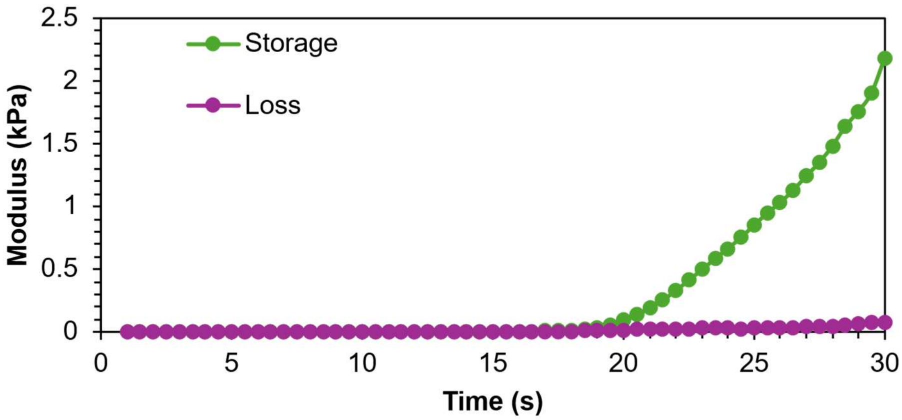


**Figure S9.** Representative NorGel photorheology data showing modulus vs. time plot to determine point of gelation. Samples consisted of 20 wt% NorGel in 80 wt% PBS, 0.2 wt% DTT, 0.3 wt% LAP, 0.1 wt% TEMPO and irradiating with a violet LED (405 nm LED, 20 mW/cm$^2$, turned on at 10 s). Sample thickness was 100 µm and samples were sealed with mineral oil to prevent evaporation.

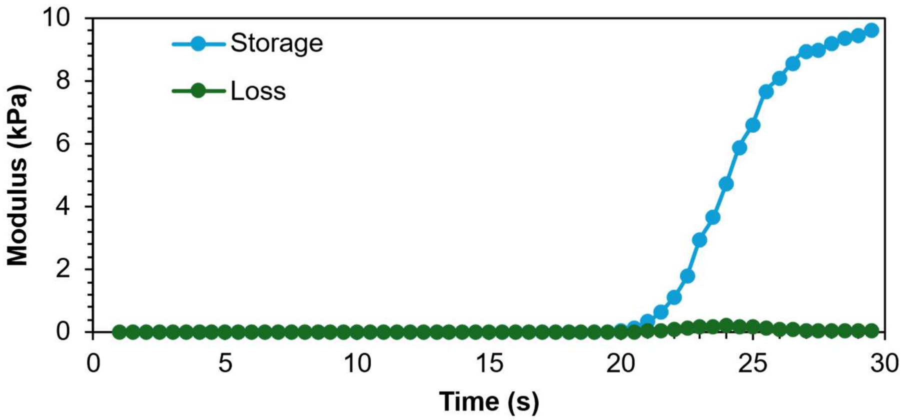


**Figure S10.** Representative NorGel photorheology data showing modulus vs. time plot to determine point of gelation. Samples consisted of 20 wt% NorGel in 80 wt% PBS, 0.2 wt% DTT, 0.3 wt% LAP, 0.1 wt% TEMPO, and irradiating with a violet LED (405 nm LED, 50 mW/cm$^2$, turned on at 10 s). Sample thickness was 100 µm and samples were sealed with mineral oil to prevent evaporation.

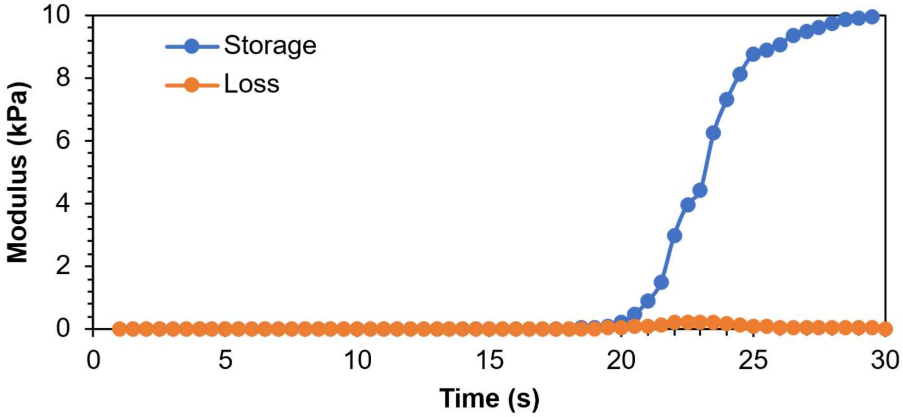


**Figure S11.** Representative NorGel photorheology data showing modulus vs. time plot to determine the point of gelation. Samples consisted of 20 wt% NorGel in 80 wt% PBS, 0.2 wt% DTT, 0.3 wt% LAP, 0.1 wt% TEMPO, and irradiating with a violet LED (405 nm LED, 70 mW/cm$^2$, turned on at 10 s). Sample thickness was 100 µm and samples were sealed with mineral oil to prevent evaporation.

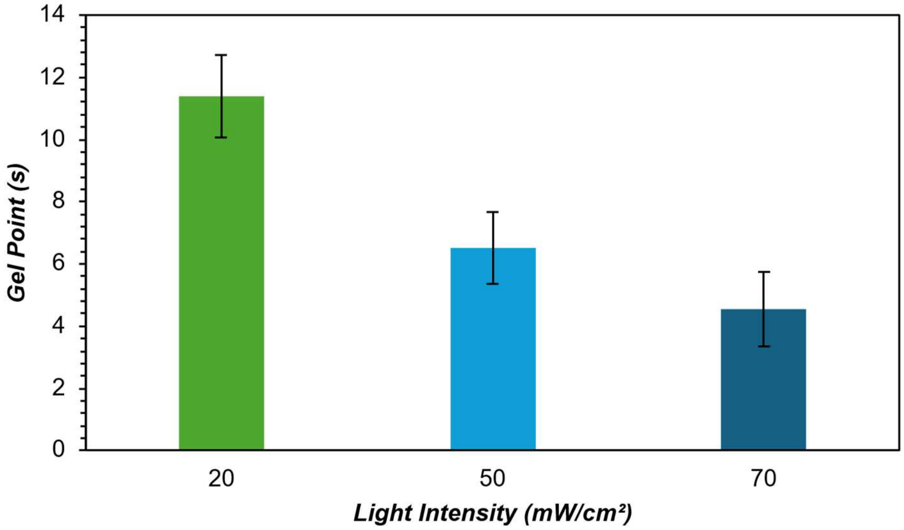


**Figure S12.** Effect of varying 405 nm light intensities (mW/cm²) on the gelation point of NorGel . Samples consisted of 20 wt% NorGel in 80 wt% PBS,0.2 wt% DTT, 0.3wt% LAP and 0.1 wt% TEMPO. Sample thickness was 100 µm and samples were sealed with mineral oil to prevent evaporation. $N = 5 \pm$ SD.

### S3. Supplementary Tables

**Table S1. Results of NorGel functionalization from different batches synthesized along with calculated amounts of dithiol (specifically, DTT) for an "ideal" (stoichiometrically balanced) step growth polymerization given a 1 g sample of 20 wt% NorGel in 80 wt% PBS.**

| *Batch* | $m_{NorGel}$ (g) | $m_{D2O}$ (g) | $mol_{TMSP}$ | $I_{TMSP}$ | *mol Nb / g NorGel* | $m_{DTT}$ (mg) |
|---|---|---|---|---|---|---|
| 1 | 0.0278 | 0.8085 | $2.35 \times 10^{-6}$ | 4.14 | $1.84 \times 10^{-4}$ | 2.8 |
| 2 | 0.0314 | 0.7391 | $2.15 \times 10^{-6}$ | 3.57 | $1.72 \times 10^{-4}$ | 2.7 |
| 3 | 0.0186 | 0.7946 | $2.31 \times 10^{-6}$ | 6.64 | $1.68 \times 10^{-4}$ | 2.6 |
| 4 | 0.0623 | 0.7762 | $2.25 \times 10^{-6}$ | 2.09 | $1.56 \times 10^{-4}$ | 2.4 |
| 5 | 0.0611 | 0.765 | $2.22 \times 10^{-6}$ | 2.43 | $1.35 \times 10^{-4}$ | 2.1 |
| 6 | 0.0212 | 0.7865 | $2.28 \times 10^{-6}$ | 6.98 | $1.39 \times 10^{-4}$ | 2.1 |
| 7 | 0.0135 | 0.7931 | $2.30 \times 10^{-6}$ | 8.01 | $1.92 \times 10^{-4}$ | 3.0 |
| 8 | 0.0361 | 0.7538 | $2.19 \times 10^{-6}$ | 3.54 | $1.54 \times 10^{-4}$ | 2.4 |
| 9 | 0.0291 | 0.7573 | $2.20 \times 10^{-6}$ | 4.55 | $1.49 \times 10^{-4}$ | 2.3 |
| | | | | **Average** | $\mathbf{1.63 \times 10^{-4} \pm 2 \times 10^{-5}}$ | **2.5** |

**Table S2. List of primary and secondary antibodies.**

| | *Antibody* | *Dilution* |
|---|---|---|
| **Primary Antibodies** | Alpha smooth muscle actin (mouse-derived) | Microscopy: 1:500<br>Flow Cytometry/FACS: 1:200 |
| | Vimentin (rabbit-derived) | Microscopy: 1:200<br>Flow Cytometry/FACS: 1:200 |
| **Secondary Antibodies** | Donkey-anti-rabbit-AF594 | Microscopy: 1:1000<br>Flow Cytometry: 1:1000 |
| | Goat-anti-mouse-AF488 | Microscopy: 1:1000<br>Flow Cytometry/FACS: 1:1000 |

**Table S3. qPCR primer sequences.**

| *Gene Name* | *Assay ID* | *Primer Sequence* | |
|---|---|---|---|
| ACTA2 | Hs.PT.56a.2542642 | Forward | 5′ AGAGTTACGAGTTGCCTGATG 3′ |
| | | Reverse | 5′ CTGTTGTAGGTGGTTTCATGGA 3′ |
| COL1A1 | Hs.PT.58.15517795 | Forward | 5′ GACATGTTCAGCTTTGTGGAC 3′ |
| | | Reverse | 5′ TTCTGTACGCAGGTGATTGG 3′ |
| COL3A1 | Hs.PT.58.4249241 | Forward | 5′ CTACTTCTCGCTCTGCTTCATC 3′ |
| | | Reverse | 5′ TTGGCATGGTTCTGGCTT 3′ |
| TBX20 | Hs.PT.58.3243363 | Forward | 5′ CAATGAACTGGATCAACATGGC 3′ |
| | | Reverse | 5′ ATGAGGCTGTGTGGTCTTTC 3′ |
| TCF21 | Hs.PT.58.20883784 | Forward | 5′ GCTACATCGCCCACTTGAG 3′ |
| | | Reverse | 5′ CACTTCTTTCAGGTCACTCTCG 3′ |

**Table S4. Storage modulus values (at 60s light exposure) for hydrogels with varying concentrations (wt% relative to bulk) of DTT. Samples consisted of 20 wt% NorGel in 80 wt% PBS, 0.1 wt% LAP and 0.1 wt% TEMPO, and irradiated with a violet LED (405 nm LED, 50 mW/cm$^2$). Sample thickness was 100 µm and samples were sealed with mineral oil to prevent evaporation. Notably, the wt% of DTT determined to be optimal (0.2 wt%) corresponded to the amount calculated for ideal stoichiometric conversion from $^1$H NMR spectroscopy. *N* = 3 ± SD.**

| ***DTT (wt%)*** | **2** | **1** | **0.6** | **0.4** | **0.2** | **0.1** | **0** |
|---|---|---|---|---|---|---|---|
| Average Storage Modulus (kPa) | 1.8 ± 0.1 | 5.0 ± 0.6 | 8.6 ± 0.4 | 12 ± 2 | 19 ± 2 | 0.25 ± <0.1 | 0.01 ± < 0.1 |

**Table S5. Effect of TEMPO concertation on NorGel point of gelation. Samples consisted of 20 wt% NorGel in 80 wt% PBS, 0.2 wt% DTT and 0.3 wt% LAP, and irradiated with a violet LED (405 nm LED, 50 mW/cm2). Sample thickness was 100 µm and samples were sealed with mineral oil to prevent evaporation. *N* = 3 ± SD.**

| ***TEMPO (wt%)*** | **0.05** | **0.1** |
|---|---|---|
| Average Gel Points (s) | 3.54 ± 0.68 | 6.69 ± 0.29 |

**Table S6. Effect of LAP concentration (wt% relative to bulk) on NorGell point of gelation . Samples consisted of 20 wt% NorGel in 80 wt% PBS, 0.2 wt% DTT and 0.1 wt% TEMPO, and irradiated with a violet LED (405 nm LED, 50 mW/cm$^2$). Sample thickness was 100 µm and samples were sealed with mineral oil to prevent evaporation. *N* = 3 ± SD.**

| ***LAP (wt%)*** | **0.1** | **0.2** | **0.3** |
|---|---|---|---|
| Average Gel Point (s) | 21.8 ± 4.9 | 13.3 ± 4.8 | 6.7 ± 0.3 |

**Table S7. Effect of varying 405 nm light intensities (mW/cm$^2$) on the gelation point of NorGel . Samples consisted of 20 wt% NorGel in 80 wt% PBS 0.2 wt% DTT, 0.3wt% LAP and 0.1 wt% TEMPO. Sample thickness was 100 µm and samples were sealed with mineral oil to prevent evaporation. *N* = 5 ± SD.**

| ***Light Intensity (mW/cm$^2$)*** | **20** | **50** | **70** |
|---|---|---|---|
| Average Gel Point | 11.38 ± 1.32 | 6.51 ± 1.17 | 4.53 ± 1.20 |
